# Fragmented micro-growth habitats present opportunities for alternative competitive outcomes

**DOI:** 10.1101/2024.01.26.577336

**Authors:** Maxime Batsch, Isaline Guex, Helena Todorov, Clara M. Heiman, Jordan Vacheron, Julia A. Vorholt, Christoph Keel, Jan Roelof van der Meer

**Affiliations:** Department of Fundamental Microbiology, University of Lausanne, CH-1015 Lausanne, Switzerland; Department of Mathematics, University of Fribourg, CH-1700 Fribourg, Switzerland; Institute for Microbiology, Swiss Federal Institute of Technology (ETH Zürich), CH-8049 Zürich, Switzerland

## Abstract

Bacteria in nature often proliferate in highly patchy environments, such as soil pores, particles, plant roots or leaves. The resulting spatial fragmentation leads to cells being constrained to smaller habitats, shared with potentially fewer other species. The effects of microhabitats on the emergence of bacterial interspecific interactions are poorly understood, but potentially important for the maintenance of diversity at a larger scale. To study this more in-depth, we contrasted paired species-growth in picoliter droplets at low population census with that in large (*macro*) population liquid suspended cultures. Four interaction scenarios were imposed by using different bacterial strain combinations and media: substrate competition, substrate independence, growth inhibition, and cell killing by tailocins. In contrast to macro-level culturing, we observed that fragmented growth in picoliter droplets in all cases yielded more variable outcomes, and even reversing the macro-level assumed interaction type in a small proportion of droplet habitats. Timelapse imaging and mathematical simulations indicated that the variable and alternative interaction outcomes are a consequence of founder cell phenotypic variation and small founder population sizes. Simulations further suggested that increased growth kinetic variation may be a crucial selectable property for slower-growing bacterial species to survive competition. Our results thus demonstrate how microhabitat fragmentation enables the proliferation of alternative interaction trajectories and contributes to the maintenance of higher species diversity under substrate competition.

## Introduction

The implications of habitat fragmentation on biodiversity constitute a widely investigated yet highly debated topic in classical macroecology ^1^. While some authors have associated fragmentation with decreasing biodiversity in general ^2^, others have reported positive effects on biodiversity at the scale of landscapes ^3,4^. In contrast, the aspect of habitat fragmentation has received only modest attention in microbial ecology ^5,6^. The formation and maintenance of taxonomically rich microbial communities in nature (*e.g.* bacteria ^7^, archaea ^8^, microbial eukaryotes ^9,10^) are thought to depend on the physicochemical conditions prevailing in their habitat ^11,12^, inherent growth characteristics of the inhabiting species ^13^, and dynamic interspecific interactions that emerge from shared nutrient and spatial niches ^14,15^. However, most of our understanding of interspecific interactions comes from experimental studies using macro-habitats, controlled growth conditions and large population census (>10^6^ cells per mL) ^13,16–18^, which are not necessarily representative for typically highly heterogenous natural microbial habitats (*e.g.* soil, plant leaves, skin). Thus, studies that take habitat heterogeneities into account are needed to extrapolate the roles of interspecific interaction effects on the development and maintenance of natural microbial communities.

Habitat heterogeneities at the scales relevant to microbial life occur in the form of spatial discontinuities and fragmentation ^19–21^, which may have important implications for microbial community assembly and diversity ^5,6,22–25^. For example, Conwill *et al.* ^26^, showed how different *Cutibacterium acnes* strains coexist at the macroscopic level of the human skin microbiome, but each individually colonizes a single skin pore; the pores creating niches without direct space and nutrient competition. Similarly, the particle structure of the soil habitat also creates multitudes of micro-pores and channels ^22,27^, which, dependent on water content, generate fluctuating networks of physically restricted and connected micro-growth environments ^20,28^. Microhabitats also arise in animal guts, because of the physical shape of the epithelial cell lining (*e.g.*, crypts ^29^), or the peristaltic motion of differently-sized food particles and mucus^30^. Surfaces of plant leaves have pronounced microstructures and properties leading to the formation of disconnected micro-droplets and water-filled channels, depending on humidity conditions ^19,31,32^. Finally, even aquatic environments, considered to be ‘connected’, are characterized by plentiful particulate organic matter to which microbes attach, forming segregated habitats ^33,34^.

Habitat space constrains the opportunities for cells to get into physical proximity to every other member of the community. Therefore, even though the diversity in a macro-environment (*e.g.*, soil) may be high (up to thousands of taxa ^7^), the spatial discontinuities lead to microhabitat fragmentation, each containing perhaps few cells from only a limited number of taxa ^27^. Communities in that regard are rather ensembles of myriads of smaller ‘sub’-communities inhabiting microhabitats. As a consequence, the assumed global roles of high-complexity interspecific interactions for community development would in individual microhabitats forcibly reduce to a few co-existing species with a smaller repertoire of potential interspecific interaction outcomes. In addition, because of the lower population census in microhabitats, one would expect phenotypic heterogeneity to play a more important role than in well-mixed conditions and at high population densities, where it would level out differences among individual founder cells. Indeed, measurements of individual phenotypic variations at the single (bacterial) cell level ^35^ show heterogeneous behaviour within low-census bacterial populations ^36–38^, and important heterogeneity in single cell growth kinetics ^39,40^. Our hypothesis here was thus that microhabitats would favour standing phenotypic variation of individual founder cells, which determines their reproductive success. In case of founder cells being part of a low-census multispecies community, we would then expect that growth kinetic variation could also lead to alternative outcomes of individual species growth and community composition. Expected interspecific interaction types from macro-scale experiments would then insufficiently explain their influence under microhabitat fragmentation. If true, growth variations in microhabitats could thus form an important driving force to sustain high observed microbial diversity despite competition for substrates being expected to drive poorly competitive species to extinction ^13,41^.

The main objective of this work was to study the effects of habitat fragmentation on bacterial community growth considering existing phenotypic heterogeneity. We tested our conjectures in four scenarios of different (expected) interspecific interaction types: (i) direct substrate competition, (ii) substrate indifference, (iii) antagonism by inhibitory compounds and (iv) direct cell killing (Fig. 1A). Strain pairs were cultured either alone or in combination, and either in standard mixed liquid suspended culture with a large starting population census to observe global interaction types, or in fragmented microhabitats with each between 1–3 founder cells (Fig. 1B). Parallel fragmented microhabitats were created by emulsifying cell suspensions in growth medium into water-in-oil picoliter-droplets (35 pL) using a microfluidic device (Fig. S1, design from Duarte *et al.* ^42^. Water-in-oil droplets have been shown previously to restrict cell movement and diffusion of compounds between droplets, and effectively shield individual growth environments ^43^. The cells in millions of generated droplets were subsequently incubated as an emulsion, and taxa growth was compared between mixed liquid suspension and pL-droplets. *Pseudomonas putida* ^44^ and *Pseudomonas veronii* ^45^ were used to test direct substrate competition and substrate independence. *Pseudomonas* Leaf 15, a known phyllosphere isolate producing inhibitory compounds ^46^ was tested with *Sphingomonas wittichi* RW1 ^47^, for antagonistic interaction effects of diffusible compounds. Finally, *Pseudomonas protegens* strains CHA0 and Pf-5 ^48^ were used to test the effect of tailocin-mediated killing. All strains were fluorescently tagged to be able to measure their productivity in individual droplets from microscopy imaging (Fig. 1C, D). Mathematical models describing competitive Monod growth ^49^ were used to examine more broadly the effects of stochastic founder cell and kinetic parameter variations on competitive strain dominance. Our results indicate that microhabitat fragmentation offers ecological opportunities to reverse interaction outcomes, leading to increased survival of poorly competitive strains in fragmented than in well-mixed bulk environments.

**Figure 1.**
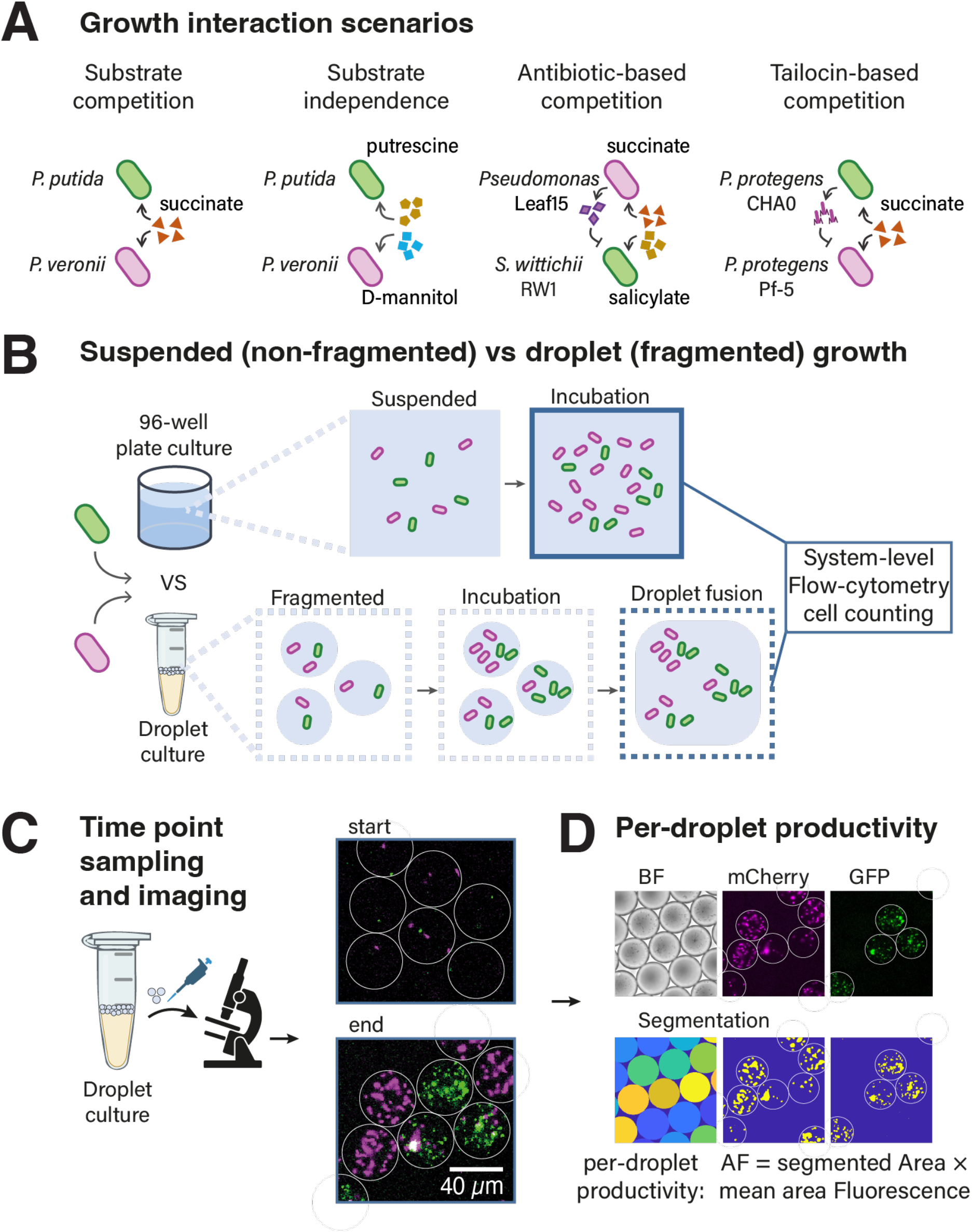
Experimental design. A) Four different pairwise interaction scenarios involving three different pairs of fluorescently tagged bacterial strains tested for growth and competition outcomes. B) Culturing in liquid suspensions in 96-well plates contrasted to that in fragmented droplets. Flow cytometry was used to count population growth. C) Droplet emulsion sampling and microscopy imaging of individual droplets. D) Productivity measurements of each strain by image segmentation of the specific fluorescent labels within segmented droplets. Reported productivity is the product of the segmented strain-specific fluorescent area (in pixels) times its mean fluorescent signal (called ‘AF’, for Area × Fluorescence).

## Results

### Growth in fragmented habitats enables local reversion of substrate competition

To understand how habitat micro-fragmentation affects the developing interactions between paired bacterial strains, we compared growth of mono- and cocultures under different interspecific interaction scenarios, and either in regular mixed liquid suspension (with a large founder population size of 5×10^5^ cells per mL) or in pL droplets (with 1–3 founder cells per droplet and strain; Fig. 1A &B, Fig. S2). In the first scenario, we focused on either substrate competition (*i.e.*, a single shared primary carbon growth substrate in form of succinate) or substrate indifference (*i.e.*, each strain receives its own specific substrate). To test this, we used two *Pseudomonas* strains (*P. putida* and *P. veronii*) with overlapping substrate preferences but different growth kinetic properties ^49^.

Growth rates of *P. putida* in liquid-suspended culture with 10 mM succinate based on fluorescence measurements (n = 6 replicates) were slightly but significantly (p = 0.0089) higher in mono than in co-culture with *P. veronii* (Fig. 2A and B). In contrast, *P. veronii* grew slightly faster in co-culture with *P. putida* than alone (p = 3.06×10^-4^). Despite this increase in co-culture, the average maximum specific growth rate (µ_max_) of *P. veronii* on succinate was 25% lower than that of *P. putida*, and the onset of growth (population lag time) took 5-6 times longer (Fig. 2A and B, Fig. S3). Consequently, liquid-suspended co-cultures became dominated by *P. putida* (Fig. 2A and C). Consistent with substrate competition, the strain-specific cell yields were lower in co-than in mono-cultures (Fig. 2C), with *P. putida* losing ca. 14.4% of its cell yield and *P. veronii* losing 84.7% compared to mono-cultures (Fig. 2C). Growth under fragmented conditions in pL-droplets both in mono- and co-cultures yielded similar cell numbers for *P. putida* in comparison to the suspended cultures (Fig. 2D, cell numbers determined after 24 h growth by flow cytometry by breaking droplet emulsions and liberating all cells into a single suspension). In contrast, the *P. veronii* cell numbers in droplet co-cultures with *P. putida* were on average 4 times higher than expected from suspended cultures (Fig. 2D). This suggested that the global competitive deficit of *P. veronii* in (macro-level) liquid suspended culture was partly abolished during growth in fragmented conditions.

**Figure 2.**
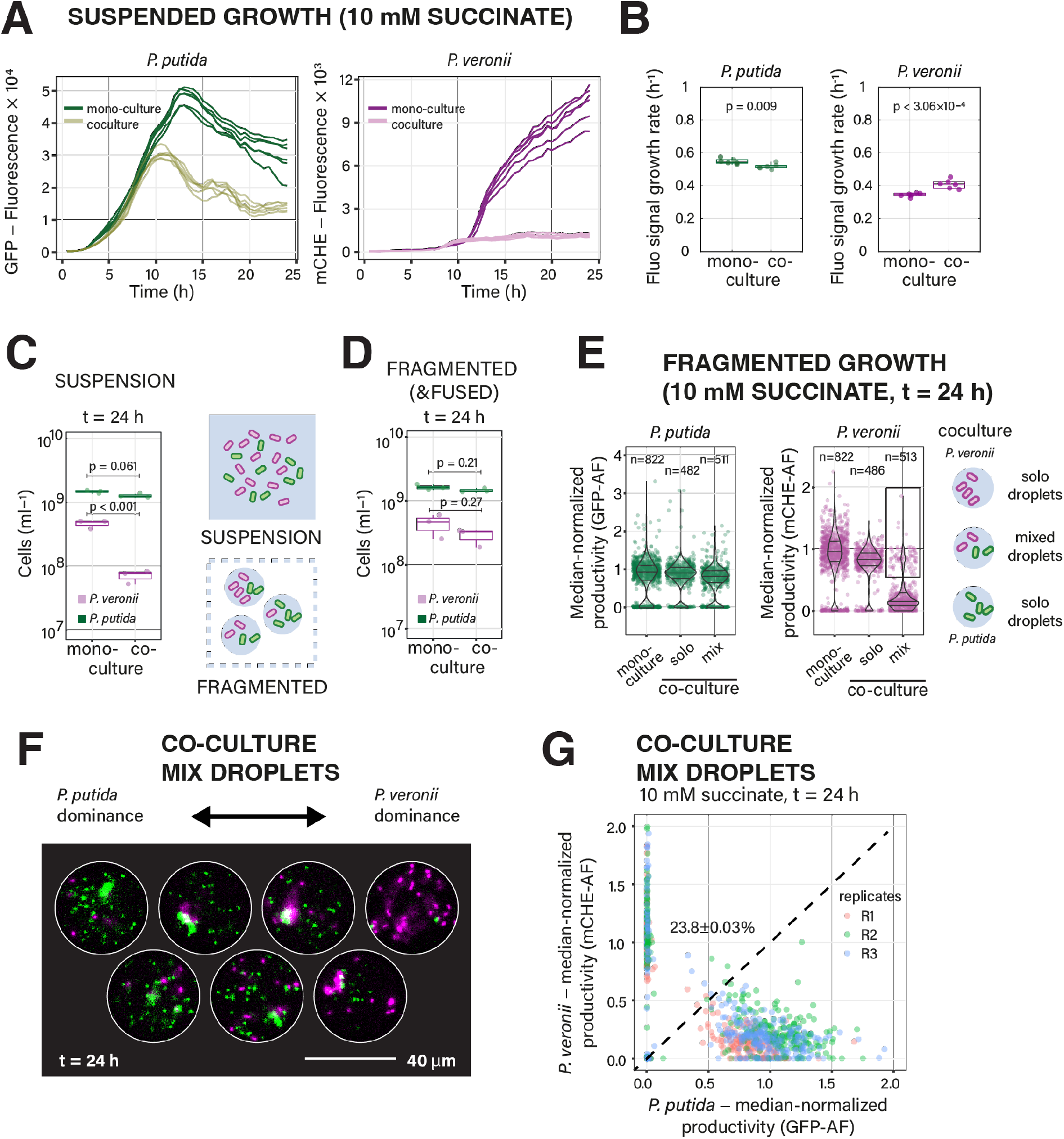
Competitive growth of *P. putida* and *P. veronii* on a single shared substrate in suspended and fragmented culture. A) Comparative growth of *P. putida* and *P. veronii* in suspended culture, alone (mono-) or in co-culture, measured by the increase of total strain-specific fluorescence. B) Inferred maximum specific population growth rates (n = 6 replicates). Plots show the median (red), lower and upper quartiles, and individual data points. P-values from two-sided t-test. C) Final population size (as cells mL^−1^) after 24 h in liquid suspension, or in broken and fused emulsions (D), measured by flow cytometry, for mono- and co-cultures of *P. putida* and *P. veronii*. Each dot is the mean count of each biological replicate (n = 3). P-values from two-sided t-tests. E) Per droplet productivity in fragmented growth conditions after 24 h (violin plot with mean, upper and lower quartiles, and individual droplet values, n = number of imaged droplets). Note the distinction between mono-culture droplets and solo droplets (*i.e.*, those droplets in a co-culture occupied by chance by only a single strain). Black frame on the right panel highlights mixed droplets where *P. veronii* reaches productivities similar to mono-cultures. F) Visual impression of the gradient of paired productivities, yielding dominance of either one of the strains. G) Median-normalized paired productivities after 24 h in droplets with both strains (n = 3 independent replicate droplet incubations). Dotted line indicates suggested equal biomass of both partners, showing dominance of *P. putida* growth but a significant proportion (23.8%) of inversed competitive outcomes (*i.e.*, *P. veronii* dominance).

To better understand the mechanisms for the attenuated competitive inhibition of *P. veronii* by *P. putida* in co-culture droplets, we looked more closely at the cell yield variations at the level of individual droplets (Fig. 2E). The median productivities of *P. putida* after 24 h growth in pL-droplets were indifferent between mono-cultures and droplets with only *P. putida* in co-cultures (Fig. 2E, *solo* droplets, p = 0.2164, n = 3), indicating that there was no difference arising from the co-culturing procedure in itself. We find such solo droplets because of the random nature of cell encapsulation in droplets, which follows a Poisson distribution (Fig. S2). Since the cell counting procedure breaks the droplet emulsion, the total counts by flow cytometry are a mixture of cells liberated from true mix droplets and solo droplets. Droplet imaging after 24 h indicated that, in co-cultures, ca. 51 % and 48 % of droplets occupied respectively by *P. putida* and *P. veronii* consist of solo droplets. The increased proportion of *P. veronii* in the fused co-culture droplet emulsions counted by flow cytometry is thus increased by the fraction of *P. veronii* solo droplets. True co-culture droplets containing both *P. putida* and *P. veronii* showed an average 11.3% reduction in median productivity of *P. putida* (Fig. 2E, *mix*, p = 0.0056 in t-test to solo droplet productivity, n = 3), which is similar as measured by flow cytometry counting on fused droplets (Fig. 2D). Productivity of *P. veronii* was indifferent between *solo* droplets (in co-culture) and *P. veronii* mono-culture droplets (Fig. 2E, p = 0.5073), but – as expected, was on average 79.4% inferior in droplets with *P. putida* present (Fig. 2E, p = 5.68×10^-4^). Mix droplet outcomes effectively ranged from those with almost exclusively *P. putida* to almost exclusively *P. veronii* and some with more equal proportions (Fig. 2F). Interestingly, in ca. 24% of co-culture droplets occupied with both strains, *P. veronii* had actually gained a competitive advantage over *P. putida* (Fig. 2G). This result indicated, therefore, that in fragmented growth conditions with low founder cell densities, *P. veronii* can overcome its general competitive disadvantage for growth on succinate.

### Fragmented growth habitats yield varying outcomes even in case of substrate independence

To contrast the reversion of competition outcomes, we next imposed a substrate ‘indifference’ scenario, in which *P. putida* and *P. veronii* were each given an exclusive carbon substrate (Fig. 1A). Our expectation here was that since substrate competition would be alleviated, both strains would grow unhindered, and liquid-suspended and pL-droplet growth would be largely similar. To test this, we used a previous observation that *P. putida* consumes putrescine but not D-mannitol, whereas *P. veronii* prefers D-mannitol and only very slowly metabolizes putrescine ^49^. Indeed, in this case, the measured growth rates in liquid suspension were indifferent between mono-and co-culture conditions for both *P. putida* and *P. veronii* (Fig. 3A, p = 0.6300, p = 0.3990, n = 6; growth curves in Fig. S4), although the time until first doubling was around 20 % shorter for *P. putida* in co-culture (Fig. 3B, p = 0.0032). Also, the total productivity (in cells mL^−1^ determined by flow cytometry) was unchanged between mono- and co-cultures, for both *P. putida* and *P. veronii* (Fig. 3C, p = 0.84, p = 0.46). In contrast, the total productivity in fragmented conditions was two-fold higher for *P. putida* than *P. veronii*, but again indifferent between mono- and co-culture conditions (Fig. 4C). Seen at population levels, these results thus suggested substrate independence for either species in liquid suspended and pL-droplet growth.

**Figure 3.**
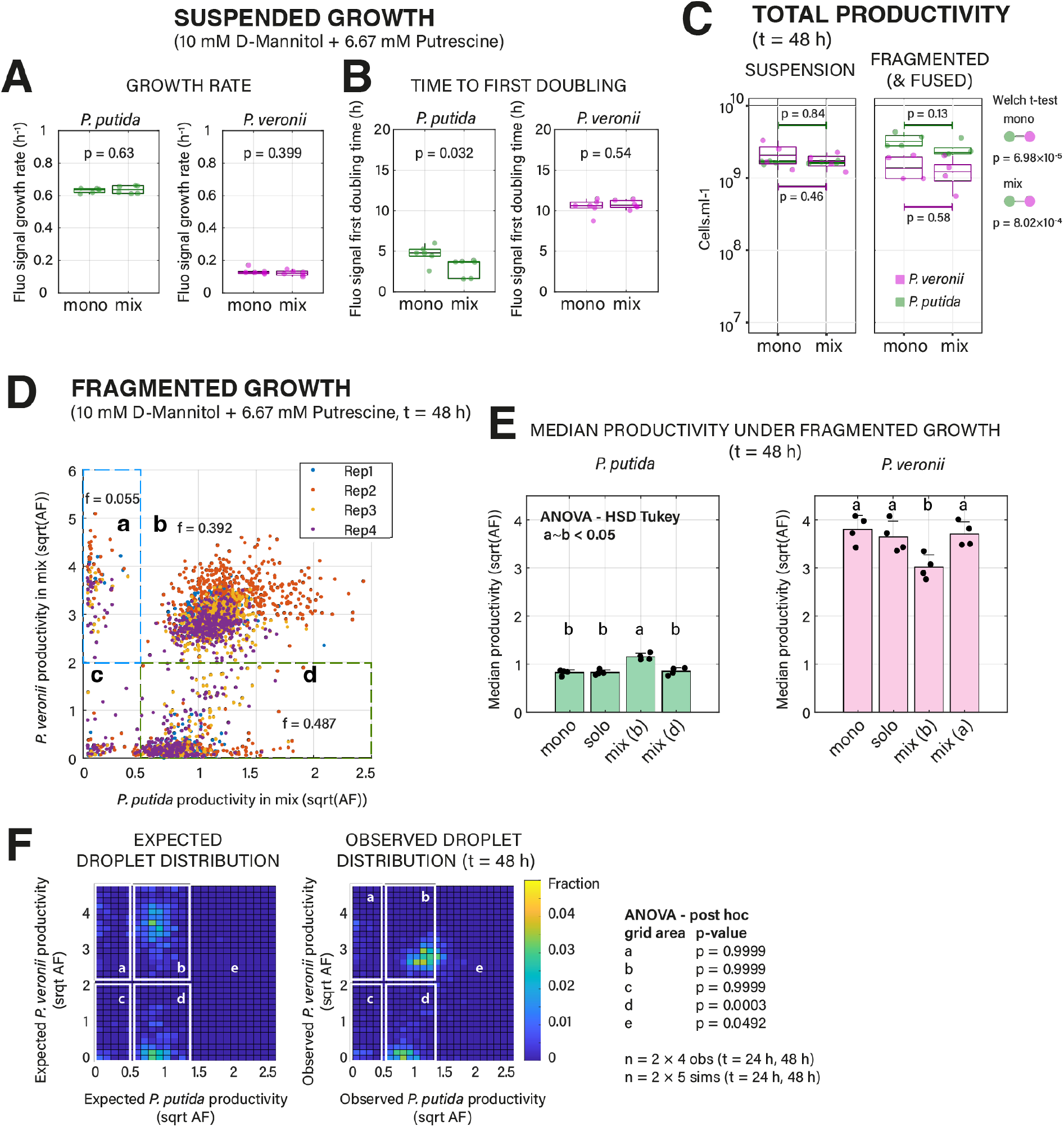
Alternative growth outcomes in fragmented growth under indifference substrate conditions. A) Maximum specific population growth rates and (B) time to first doubling of mono- and co-cultures of *P. putida* and *P. veronii*, measured in 96-well plate liquid suspension in the presence of two substrates (prioritized by either of the strains: D-mannitol for *P. veronii* and putrescine for *P. putida*; n = 6 replicate values summarized by box plots with individual data points, p-value from two-sided t-test; growth curves in Fig. S4). C) Total population size (as cells mL^−1^) measured by flow cytometry after 48 h for mono- and co-cultures of *P. putida* and *P. veronii* in liquid suspension (left panel) or in droplets (fused emulsions, right panel). Each dot is the mean count for each biological replicate (n = 4). D) Paired productivities after 48 h of *P. putida* and *P. veronii* in mixed droplets, overlaid from four independent biological replicates (REP1–4, individual colors), cultured on D-mannitol and putrescine (square-root-transformed AF-values). Fractions (*f*) denote the mean values of the proportion of paired droplet productivities falling in the grid areas labeled **a–d** (separated by dotted lines). E) Increased median productivities for *P. putida* (left panel) but not for *P. veronii* (right panel) in mix droplets area **b**, compared to mono, solo and mix-area **d** droplets. Statistical differences between median productivities (n = 4 biological replicates) in mono and co-culture droplets were determined with ANOVA coupled with HSD Tukey test, and are indicated with letters a and b. F) *P. putida* grows better than expected in mix droplets with *P. veronii* (seen here as shift into grid area **e**). p-values from ANOVA post hoc multiple test, with n = 4 biological replicates and n = 5 simulations, combined from two time points). Expected droplet distribution for pairs simulated from individual mono-culture droplet distributions at the same time point (assuming no interactions).

**Figure 4.**
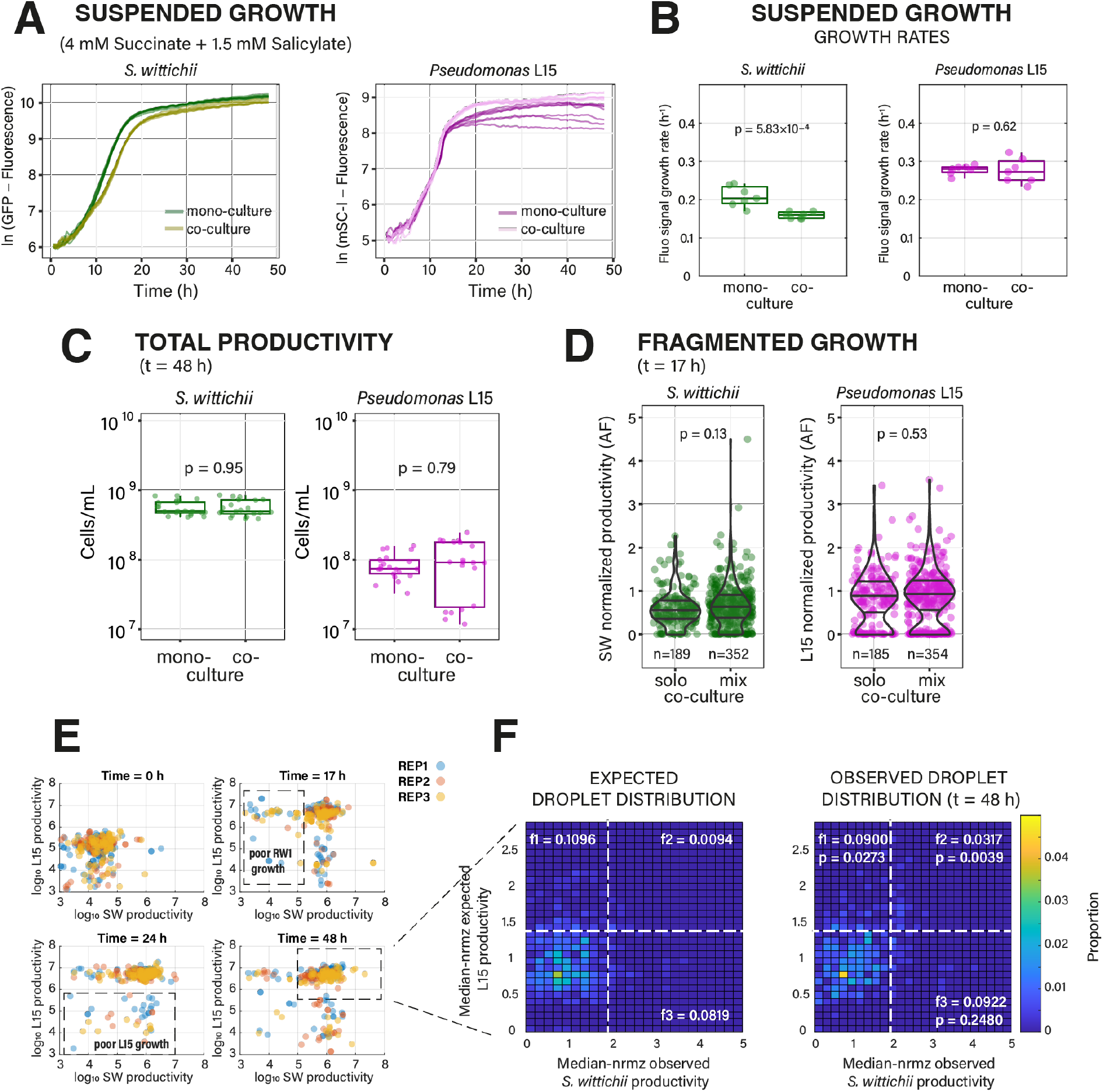
Fragmented growth can lead to overturning of inhibition under indifference substrate conditions. A) Growth of *S. wittichii* RW1 (SW) and *Pseudomonas* Leaf15 (L15) cultured separately or together in 96 well-plates under substrate independence (n = 7, measured from strain-specific fluorescence). For growth on individual substrates, see Fig. S5. B) Maximum specific population growth rates in suspended growth conditions (n = 7). P-values from Wilcoxon rank-sum tests. C) Total productivity (as cells mL^−1^) measured by flow cytometry after 48 h in suspended culture (n = 3 biological replicates with each 7 technical replicates). P-values from two-sided t-tests between mean counts of biological replicates. D) Per droplet productivities of either strain in solo and mix fragmented culture at t = 17 h (n = number of observed droplets; normalized to the median of the solo droplets at t = 48 h). P-values from Wilcoxon rank sum tests. E) Strain-specific productivities over time in mix droplets (log_10_-transformed AF values, three replicates overlaid, each subsampled to 100 droplets), indicating the ca. 10 % of droplets with poor growth of either partner. F) Comparison of expected vs observed growth of pairs, removing the poorly-growing fractions (i.e., enlarged boxed region in panel E). AF-values normalized by their distribution maximum (roughly equivalent to the median). Paired distributions arbitrarily separated to fractions above- or below average productivity properties to facilitate comparison (n = 5 simulations for each time point and observation replicate, n = 3 replicate observations, three time points). P-values from Wilcoxon sign-rank test including the same grid fraction across all replicates and time points (*f* denotes their mean). Expected paired droplet distributions simulated from observed solo droplet growth (assuming no interactions).

At the level of individual droplets, the substrate indifference scenario presented itself again very differently. An average of 5.5 % of droplets were dominated by *P. veronii* (Fig. 3D, fraction **a**), whereas 39.2 % consisted of droplets where productivities were equal (Fig. 3D, fraction **b**, n = 4 biological replicates). In contrast, 48.7 % of co-culture droplets (*i.e.*, having detectable fluorescence signals of both *P. putida* and *P. veronii*) were largely dominated by *P. putida* (Fig. 3D, fraction **d**). The median productivity of *P. putida* was higher in the fraction **b** droplets (Fig. 3E, ANOVA, post-hoc p = 0.0213, compared to *P. putida* solo droplets), whereas that of *P. veronii* in fraction **b** was lower compared to *P. veronii* mono-culture droplets (Fig. 3E, p = 5.67 × 10^−4^; n = 4 replicates). Compared to a *null* model of co-culture droplet distributions, the productivity of *P. putida* was indeed significantly higher in mix droplets with *P. veronii*, but significantly lower when being in droplets alone, than expected from the sampled probability of its individual productivities in mono-cultures (Fig. 3F, ANOVA with post-hoc test, grid fractions **e** and **d**, respectively). This indicated that interactions in droplets with equal-sized partner populations were mutualistic for *P. putida* and slightly antagonistic for *P. veronii*. These results thus illustrated how a globally perceived non-competitive scenario breaks down in a variety of different outcomes in a fragmented habitat.

### Fragmented growth effects under an inhibition scenario

To explore whether fragmented growth would more generally result in more variable outcomes than in mixed liquid culture, we used two other strain combinations and highlighting biological interactions other than substrate competition.

In the first of these, we produced an inhibition scenario, consisting of a phyllosphere isolate *Pseudomonas* sp. Leaf15, known to excrete a growth-inhibitory compound ^46^, mixed with a sensitive strain (for which we used a fluorescently tagged variant of *S. wittichii* RW1 ^47^). In this scenario both strains have their own carbon substrate, to avoid generating additional substrate competition (Fig. 1A). We used succinate for L15, which is not measurably used by RW1, and salicylate for RW1, which is not used by nor toxic for L15 (Fig. S5). As expected, growth rates of RW1 in co-culture liquid suspension with L15 were reduced by 25% compared to its mono-culture, whereas those of L15 are unaffected, confirming growth rate inhibition (Fig. 4A, 4B). Despite the growth rate decrease, the final attained population size of both RW1 and L15 in liquid suspension co-culture was indifferent from the mono-cultures (Fig. 4C, measured by flow cytometry; p = 0.95, p = 0.79). Also, the productivity of RW1 in mix droplets with L15 was similar to that in solo droplets (Fig. 4D, p = 0.13, p = 0.53, n = 4; Fig. S6), although both showed a constant ca. 10 % fraction of non- or poorly growing cells (Fig. 4E). Compared to mono-culture growth, productivity of RW1 was the same and that of L15 slightly higher in co-culture mix droplets (p = 0.0020; signrank test on median of the growing droplet fraction, Fig. 4F, Table S1). However, there was a 0.8–9.3 % (average 3.2 %) fraction of mix droplets with RW1 and L15 productivity higher than expected from their mono-culture droplet growth (Fig. 4F, *f2* fraction, p = 0.0039, signrank test all time points and replicates) reached between (Fig. 4E, *f*2 fraction, three time points). This fraction thus represents local positive interactions, suggesting reversal of inhibition under fragmented growth conditions.

### Tailocins provide a competitive advantage only within fragmented habitats

In the final example, we studied the interactions between two *P. protegens* strains, one of which (Pf-5) is sensitive to a phage tail-like weapon, or tailocin, produced by the other (CHA0), leading to its lysis ^48^ (Fig. 1A). CHA0 is self-resistant to its own tailocins ^48^. Activation of tailocin production and release, however, is a rare event in CHA0 cultures and requires a stress trigger ^48^. Consequently, we expected that variable tailocin production may occasionally change the competitive outcome during growth on the same substrate, which would be detectable under fragmented growth, but not in mixed liquid suspended cultures.

Co-cultured strains on a single common substrate (succinate) in liquid suspension indeed yielded almost equivalent substrate competition outcomes, with equal time to reach stationary phase for both CHA0 and Pf-5 in mono-cultures, and ca. 50/50 yields in stationary phase (Fig. 5A). Productivities of either strain in co-culture pL-droplets were also equal and approximately half of that in mono-culture droplets (Fig. 5B, *solo*). The observed distribution of the productivities of Pf-5 and CHA0-in mix droplets followed an almost perfect constant ‘sum’, composed of the variation of individual productivities of Pf-5 and CHA0 (Fig. 5C).

**Figure 5.**
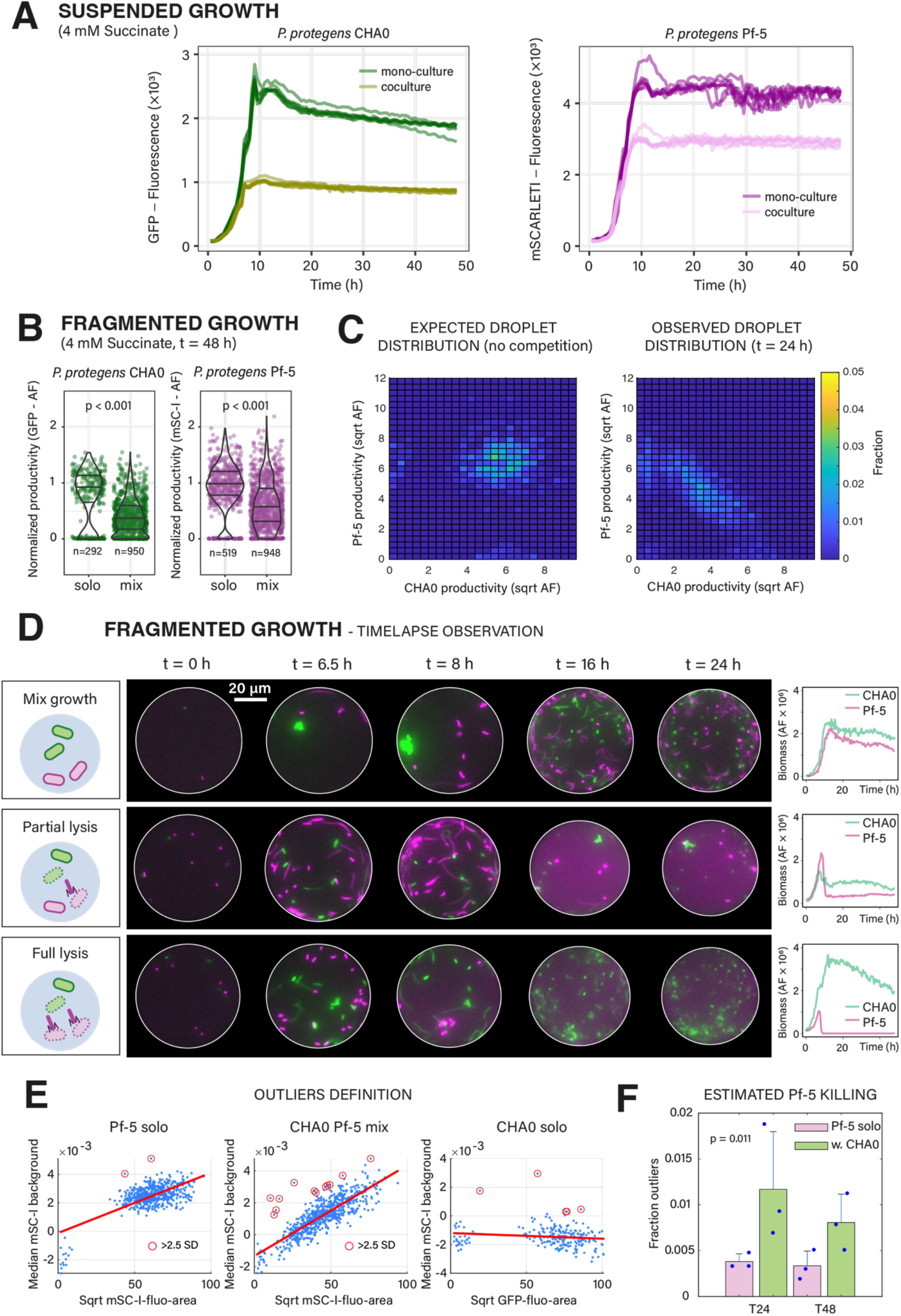
Heterogeneous tailocin production ensures infrequent competitive dominance in microhabitats. A) Equivalent growth of *P. protegens* CHA0 and Pf-5 cultivated separately or together with 4 mM succinate in liquid suspension (n = 7 replicates). B) Per droplet productivities of both strains in solo and mix droplets after 48 h (normalized to the median of the solo droplets). P-values from Wilcoxon rank-sum tests. C) Expected (from solo droplets, without competition) and observed mix droplet productivities after 24 h, indicating almost equal competitive substrate sharing (square-root transformed AF-values). D) Effect of tailocin lysis in individual droplets. Top: mixed growth without lysis; middle, partial lysis of Pf-5 (in magenta); bottom, total lysis of Pf-5 in the presence of CHA0. Time points selected from time-lapse imaging. Strain-specific fluorescence (biomass) development shown on the right. E) Variation in mScarlet-I (Pf-5 solo droplets) fluorescent background sets outlier range, above which Pf-5 lysis is assumed (red-circled data points, > 2.5 times standard deviation of the residual variation to the linear regression line). F) Inferred fractions of Pf-5 lysis in the presence of CHA0 compared to Pf-5 solo background (n = 3 replicates, p-value from two-sided *t*-test on combined t = 24 h and t = 48 h outlier fractions).

Interestingly, however, in a small fraction of individual droplets with both CHA0 and Pf-5, an increased background fluorescence in the mScarlet-I fluorescence channel for Pf-5 could be observed, which in timelapse droplet imaging appeared as sudden onsets of partial and even complete disappearance of Pf-5 cells (Fig. 5D, Supplementary movie S1). This sudden disappearance of Pf-5 cells would be in agreement with the release of tailocins from CHA0 leading to the puncturing and liberation of the cell content of the sensitive Pf-5 cells (consequently leading to an elevated background fluorescence by diffusion of mScarlet-I protein). From the variation of Pf-5 median background fluorescence in solo droplets (Fig. 5E), we estimated that on average ca. 0.5% of all droplets with both partners show evidence for lysis of Pf-5 (*i.e.*, above 2.5 × the Pf-5 solo background standard deviation, Fig. 5E; Fig. 5F, p = 0.0114, n = 3 biological replicates, two time points combined). In summary, these results indicated that both *P. protegens* strains are equally competitive for succinate, but that the production of tailocins by CHA0 can help to remove the competitor. Tailocins can thus have a crucial localized effect in co-inhabited microhabitats, but this effect is masked in liquid-suspended culture, because of their low activation rate.

### Phenotypic variation in growth kinetics of founder cells determines colonization outcomes in microhabitats

Since all the co-culture outcomes under fragmented conditions showed important variability compared to well-mixed bulk conditions, we wondered if this would be the result of inherent founder cell phenotypic variability. To demonstrate this, we turned our attention again to the case of *P. putida* and *P. veronii* and a single competitive substrate, and measured growth in individual droplets over time. For this, we used microfluidic chips with a low ceiling (10 µm height) ^50^, so that droplets are squeezed, kept in place and better cell focusing is obtained. Although the incubation in PDMS-glass results in slightly different oxygen provision to growing cells than culturing them in a pL-droplet emulsion, it enabled measuring the variability of cell growth in individual droplets (Fig. 6A). Indeed, timelapse imaging confirmed different outcomes from the same starting configurations (*e.g.*, one *P. putida* cell and one *P. veronii*, Fig. 6B, at t = 0 h), and growth measurements of n = 191 individual droplets showed kinetic variability both in mono- and co-culture droplets (Fig. 6C). Average growth rates of *P. putida* in solo droplets were 1.2 times higher than in mixture, whereas those of *P. veronii* remained indifferent between the two conditions (Fig. 6D). On average, *P. veronii* started dividing 4 h later than *P. putida* (Fig. 6E). Both strains also showed a tendency that incidental longer lag times decreased their final attained size in co-culture droplets (Fig. S7). Paired growth trajectories were highly variable between droplets, even under the same starting cell-census (Fig. 6F and Fig. S8), whereas unequal starting cell ratios tended to favour either one of the strains (Fig. 6G). However, growth rates and lag times of *P. veronii* were the only significant predictors for biomass ratio outcomes (generalized linear mixed effects model, r^2^ = 0.8236, n = 108 co-culture droplet pairs, Fig. 6H, Table S2), whereas founder cell numbers were less relevant (Table S2). The variance in single droplet growth rates and lag times tended to decrease with increasing starting cell numbers (Fig. 6I, Fig. 6J, significant inequality of variances for *P. putida* but not for *P. veronii* - Brown-Forsythe test, see parameter distributions in Fig. S9), suggesting that the influence of individual cell heterogeneities becomes less determinant and yields more averaged behaviour.

**Figure 6.**
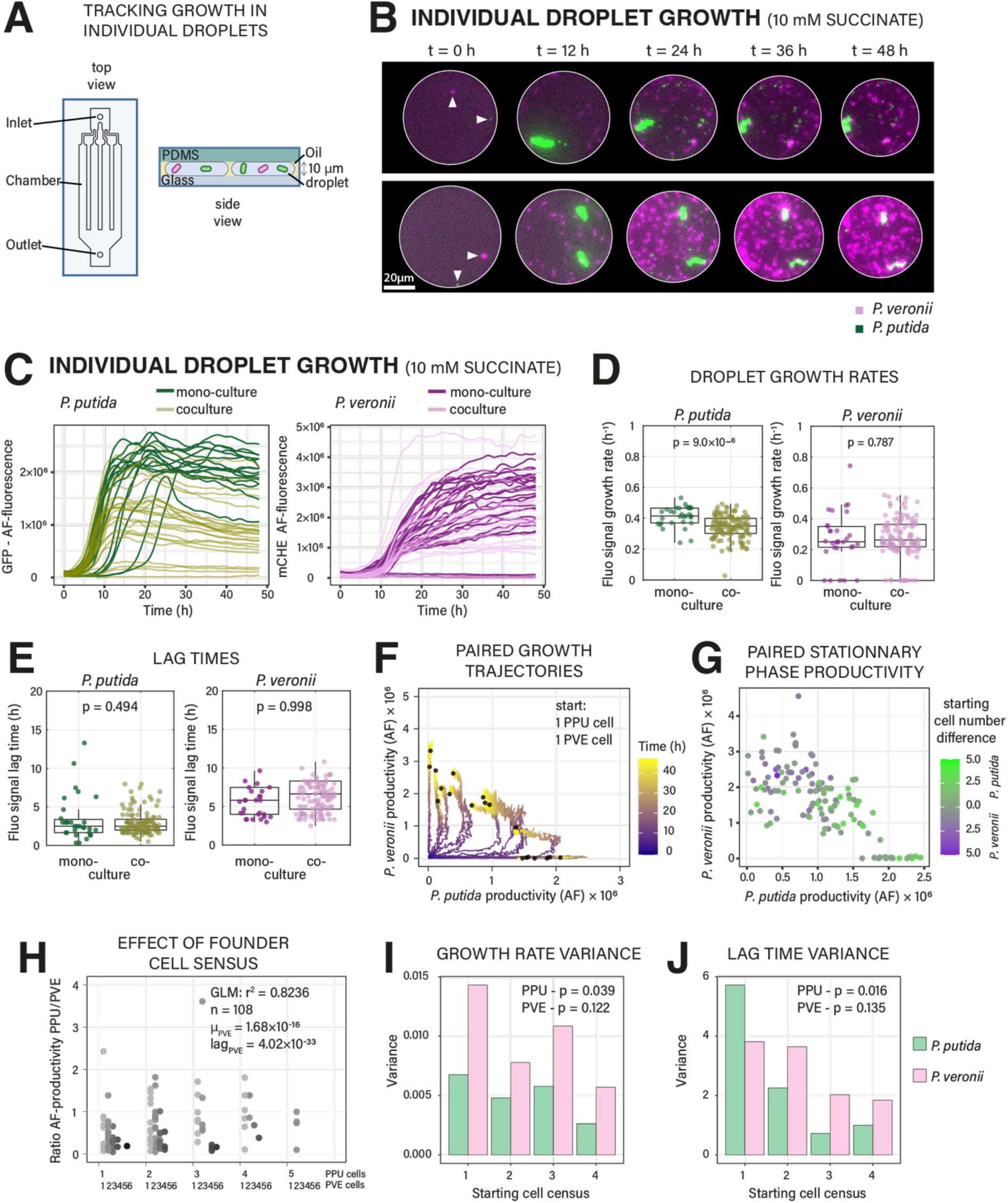
Heterogeneous growth kinetics and variable founder cell number determine competitive growth outcomes. A) Schematic of the observation chip used to track population growth within single droplets over time; figure panel adapted from Ref. ^50^. B) Time series examples of two droplets with the same number of founder cells for both *P. putida* and *P. veronii*, but with different outcomes. C) Growth in individual droplets of either strain in mono- or co-culture. D) Inferred maximum specific population growth rates and lag times (E) in individual droplets of panel C. Averaged data is represented in box plots with individual data points shown on top. P-values from two-sided *t*-testing. F) Different growth trajectories of pairs with both a single founder cell, showing productivity of each member as its specific fluorescence (non-transformed AF-values). Black dots indicate final AF-productivities (t = 48h). G) Stationary phase per-strain productivity in mix droplets as a function of founder cell number (colours showing the difference in the number of starting cells, towards *P. putida* majority in green or *P. veronii* majority in magenta). H) Effect of founder cell census on the ratio of stationary phase productivities (mean stationary AF-values) of *P. putida* (PPU) and *P. veronii* (PVE) in mix droplets. The initial number of both *P. putida* and *P. veronii* cells is indicated on the x-axis. Variation in the productivity ratio is best explained by variations in growth rates (µ_PVE_) and lag times (lag_PVE_) of *P. veronii* (General linearized mixed effects model, r^2^ = 0.8236, see Table S2). Effect of founder cell census on the variance in the measured maximum population growth rates (I) and lag times (J) of either strain in droplets (mono and mix combined). P-values from Brown-Forsythe test.

To better demonstrate the effect of single-cell growth variation on competitive outcomes in a two-species community within the fragmented habitat, we adapted an existing mathematical framework ^49^ for simulating carbon-limited competitive Monod growth of *P. putida* and *P. veronii* founder cells within 35 pL-droplets (Fig. 7A). In this simulation, each founder cell starts with independent growth kinetic parameters, subsampled from inferred distributions around means measured in liquid mono-cultures (Fig. 7A). In addition, each droplet is colonized by a Poisson-random number of founder cells. Simulations including both individual growth kinetic variability and Poisson-distributed initial cell ratios (λ = 3 for both *P. putida* and *P. veronii*) produced growth distributions in co-culture droplets consistent with the observed distributions of *P. putida* and *P. veronii* co-culture droplet productivities (Fig. 7B, 7C). In contrast, both a uniform starting distribution or uniform individual growth kinetics reduced the simulated fraction of droplets in which *P. veronii* reverses competition (Fig. 7D, 7E). Further simulations suggested then that mostly an increased heterogeneity of lag times and growth rates of *P. veronii* founder cells determines reversed competition outcomes with *P. putida* (Fig. 7F–7H). This agrees with them being the most important predictors in the general linear mixed model outcome (Fig. 6H). Increasing kinetic heterogeneities among *P. putida* founder cells are also predicted to positively impact the reversal of competition by *P. veronii*, suggesting that *P. veronii* benefits from the heterogeneity among its competitor cells (Fig. 7F– 7H). Collectively, these results and simulations underscore that heterogeneity in growth properties among founder cells can enhance the probability of variable outcomes between competing species inhabiting fragmented habitats. This may also imply that some bacterial species have become selected for more variable growth kinetics, which can favour their survival under substrate competition in microhabitats.

**Figure 7.**
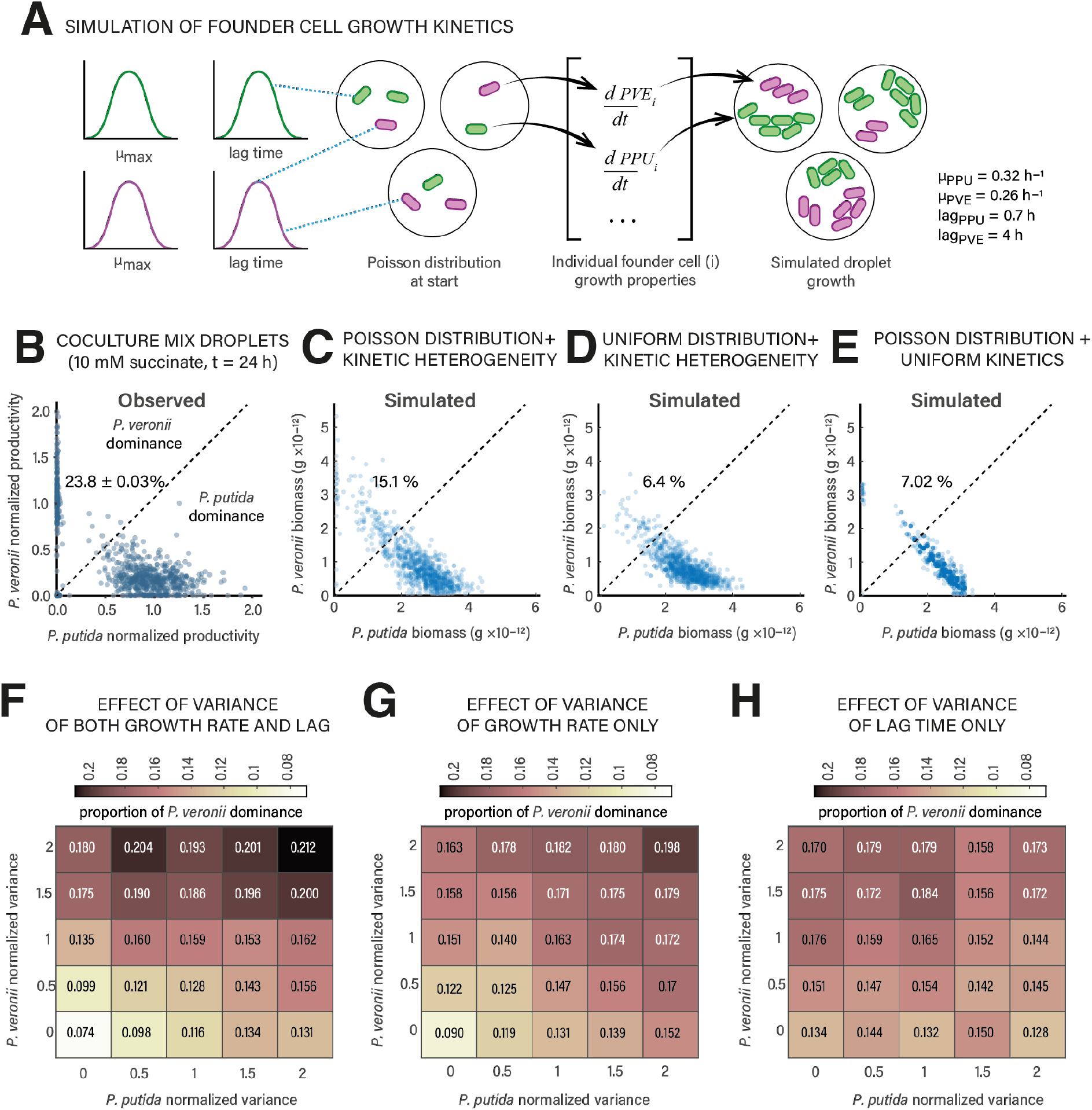
Model simulations predict a positive effect of founder cell kinetic heterogeneities on competitive co-existence in fragmented habitats. A) Model workflow to simulate Monod substrate-limited (10 mM succinate) competitive growth of *P. putida* (PPU) and *P. veronii* (PVE) in droplets from Poisson-picked individual founder cells, with individual subsampled growth rates and lag times (average values inferred from liquid suspended mono-culture growth, as in Fig. S3). For model details, see methods section. B) Observed strain productivities of *P. putida* and *P. veronii* in mix droplets on 10 mM succinate after 24 h (reproduced from Fig. 2G for ease of comparison; normalized by the median strain-specific fluorescence of the droplet mono-cultures), in comparison to simulations with either both Poisson-distributed starting conditions and kinetic heterogeneity (C), only kinetic heterogeneity (D) or only Poisson-distributed founder cells (E). Percentages indicate the proportion of droplets with dominant *P. veronii* growth (*i.e.*, either normalized fluorescence or simulated biomass above the diagonal trend line). Dots represent individual observed or simulated droplets (n = 3 replicates or simulations, 1000 droplets each). F) – H) Effects of variance in growth kinetic parameters of either strain on the proportion of simulated *P. veronii* dominance, either combined (growth rate and lag time) or individually. Normalized variance of 1 corresponds to the parameter values used in simulations of panel C.

## Discussion

Here we demonstrate how phenotypic variability in single-cell growth kinetic parameters and Poisson-variation in assembly of strain pairs in micro-scale communities lead to distinct ecological outcomes and overturning of global-scale inferred interaction types. We show, using three different strain pairs and four different imposed growth regimes and interactions, how growth in fragmented environments enables local competitive overturning, whereas culturing at large population census does not. This aspect of interaction outcome reversal has received little attention in previous studies describing paired species interactions. Even though the fraction of interaction reversals may seem relatively small, this effect can help to explain why less competitive species can locally sustain in mixed microbial communities within fragmented environments ^51^.

An important premise for our work was to consider that natural environments for microbial communities are characterized by a high degree of spatial fragmentation and/or compartmentalization. Secondly, we assumed that such fragmentation and compartmentalization occur at a relevant micro-scale, such that the formed microhabitats are colonized by low numbers of founder cells and species. There is plentiful evidence to support the assumption that local habitats for prokaryotic cells measure in micrometer dimensions with low population census ^22,52^. For example, an estimated 90% of microbial cell clusters in soils contain fewer than 100 cells ^28^; plant surface architecture is characterized by micrometer crevices and microdroplets enabling microcolony formation ^19,31^, and sinking food particles in the ocean range in sizes from 1–50 µm with initial colonization numbers of 10^2^-10^3^ cells ^53^. In addition, cell-cell interactions are assumed to be dominating at short (10-100 µm) distances ^54^. Therefore, microscale habitats with low numbers of founder cells potentially amplify any standing phenotypic variation, which will control the success of their local proliferation.

To measure the presumed effects of fragmentation on reproductive success, we relied on microfluidic pL-droplet formation and cell encapsulation. Droplet cultivating approaches have attracted interest as a high-throughput method to co-culture bacteria ^15,36,37,50^, or enrich bacteria from natural samples in an untargeted manner ^55,56^, and potentially allowing coculturing of unculturable members in multi-species conditions ^57^. We adopted the method here to study variability in growth and assembly of paired bacterial populations under differently imposed growth regimes and interaction types, which can mimic the events that may occur in natural fragmented environments. Notably, the droplet encapsulation creates isolated habitats that only allow local resource depletion and the development of metabolic or contact-dependent interactions, but no cross-talk between droplets ^43^. In contrast to regular liquid suspended cultures with larger volumes (here: 140 µL) and starting cell densities (10^6^ cells), which confirmed the intended global interaction types (*i.e.*, substrate competition, indifference, and growth rate inhibition), we measured persistent effects of interaction reversal under fragmented growth conditions. For example, *P. veronii* grew to 4 times higher densities in fragmented co-culture droplets than under the same competition with *P. putida* in liquid culture, and ca. 24% of mix droplets became dominated with *P. veronii* (Fig. 2). Global substrate indifference, in contrast, led to the opposite: the appearance of individual droplets with higher-than-expected growth of either of the partners (Fig. 3). Global growth rate reduction on *S. wittichii* RW1 by *Pseudomonas sp.* Leaf15 was also overturned with 1-9 % of isolated mix droplets having higher than expected growth of the sensitive partner (Fig. 4). Timelapse imaging and mathematical simulations demonstrated that it is likely that the competitive overturning is due to inherent physiological differences among founder cells, as we hypothesized. Given that bacterial cell-to-cell phenotypic variation is a general phenomenon ^35–37,50^, we can thus reasonably extrapolate our findings to most paired and higher-order communities starting from small numbers of founder cells from different taxa, and assume that microscale habitat fragmentation in general, would lead to more variable outcomes than expected from globally observed interaction types.

Our results do not only hold for interactions mediated by metabolic products but also for interactions involving bacterial killing by specialized weapons such as tailocins. Fragmented growth in isolated droplets showed that, despite low production rates, tailocin production by *P. protegens* CHA0 can eradicate *P. protegens* Pf-5 in microhabitats, whereas this killing has no effect in bulk liquid cultures (Fig. 5). The production of tailocins is indeed a highly heterogeneous process at a population level, initiated in less than 1% of cells ^48,58^. In addition, tailocins, like other specialized bacterial killing devices such as the type VI secretion system, act very locally ^48^, which has raised the question of their ecological importance ^58^. Our droplet observations suggest that tailocins are particularly helpful in overcoming competition in spatially restricted microhabitats. Within the context of the natural habitat of *P. protegens* (the plant rhizosphere), killing by tailocins could help to maintain local ‘reservoirs’ of the producer strain, possibly increasing its survival and ability to colonize new habitats.

How important are our findings for explaining the general high diversity of microbial communities? One of the larger conundrums in microbial ecology is explaining why so many taxa with overlapping metabolic capacities but different growth rates can co-occur in the same habitat, as one would expect the slowest growers to go extinct as a result of competition ^13,41^. One can argue that slight differences in substrate utilization and metabolic dependencies may provide opportunities for co-existence ^59^. In addition, the (dynamic) spatial fragmentation of habitats can contribute to co-existence ^20,60^. Indeed, several reports show that taxa diversity prevails even at small scales, suggesting effects of spatial isolation and colonization bottlenecks. For example, individual pores on human skin are colonized by different strains of *C. acnes* ^26^ and bacterial community composition within soil micro-aggregates is highly variable, even below 20-µm particle size ^10,61^. However, spatial fragmentation itself is not enough to maintain diversity if taxa would not show phenotypic variation, because, in the absence of cell-cell variability, interspecific interactions would become completely deterministic. We thus conclude that the importance of the micro-scale is not simply to provide spatial isolation, but to integrate variations in local interspecific interactions and founder cell numbers, which then drive the emergent composition of the local community under growth. Spatial fragmentation (or perhaps rather: dynamic variation in spatial fragmentation) thus plays a crucial role in types of local interactions and the resulting diversity of a complex meta-community ^5,6^. To extrapolate downwards from globally measured interactions to small scales is not doing justice to the existing variability in such interactions and provides an oversimplification of their role in community development.

## Material and Methods

### Strains, media and culture conditions

Two *Pseudomonas* strains were used for the substrate interaction experiments: *P. putida* F1 (PPU) is a benzene-, ethylbenzene- and toluene-degrading bacterium from a polluted creek ^44^. *P. putida* F1 was tagged with a single-copy chromosomally inserted mini-Tn5 cassette carrying a constitutively expressed fusion of *eGFP* to the P_circ_ promoter of ICE*clc* ^62^. *P. veronii* 1YdBTEX2 (PVE), a BTEX-degrading strain, was isolated from contaminated soil in the Czech Republic ^45^. PVE was tagged with a constitutively expressed *mCherry* from the P_tac_-promoter within a single-copy inserted mini-Tn7 transposon (carrying a P_tac_–*mCherry* cassette - as described in ^63^). For the growth inhibition experiment, we used *Pseudomonas sp* Leaf15 (L15), an antibiotic-producer isolated from *Arabidopsis thaliana*’s phyllosphere ^46^, and a fluorescently tagged version of *Sphingomonas wittichii* RW1 ^47^. L15 was tagged with a constitutively expressed *mScarlet-I* single-gene copy, using a pMRE-Tn7-145 mScarlet-I plasmid ^64^. We used two rhizospheres inhabiting strains of *Pseudomonas protegens*, CHA0 and Pf-5, for the tailocin interaction experiment. CHA0 was tagged with a constitutively expressed single inserted gene copy of *gfp2* ^48^, and Pf-5 was tagged with a constitutively expressed *mScarlet-I* from a single-copy inserted mini-Tn7 transposon (using a pUC18T-mini-Tn7T-Gm-Pc-mScarletI plasmid).

Strains were streaked on a nutrient agar plate directly from a -80°C stock and were incubated for 2-3 days at 30°C before being stored at 4°C for the later experiments (max 12 days). Each biological replicate was started from a single isolated colony of each strain, which was resuspended in a Mc-Cartney glass tube with 5 mL (or 8 mL for L15 and RW1) of 21C minimal medium (21C MM, as described by Gerhard *et al.* ^65^) supplemented with the appropriate carbon substrate(s) (see Table S3). Cultures were incubated at 30°C under rotary shaking at 180 rpm.

Precultures were centrifuged in 50-mL Falcon tubes to harvest the bacterial cells. The cells were two times successively washed in 5 mL of 21C MM before being resuspended by pipetting in 5 mL of 21C MM. *P. putida* and *P. veronii* cultures were centrifuged for 4 min at 12,000 rpm (Eppendorf centrifuge 5810R with an F-34-6-38 rotor, 6×15/50mL conical tubes), whereas the other four strains were centrifuged for 4 min at 8,000 rpm. Specifically, the *S. wittichii* suspension was vortexed for 2 min at the final resuspension step to disperse cell aggregates as much as possible. The turbidity of the final cell suspension was measured with a spectrophotometer (MN – Nanocolor Vis, OD_600_), and then diluted in 21C MM with the appropriate C-source(s) to have approximately the same starting cell numbers (OD_600_ of 0.02– 0.05, depending on the strain; see Table S3). For co-culture experiments, the diluted cell suspensions of the respective partner strains (Table S4) were mixed at a 1:1 ratio (*vol*/*vol*). Mono-culture controls were diluted two times with 21C MM including the appropriate C-source(s), thus maintaining the same starting density for each strain in mono- and co-culture.

### Liquid suspended growth in 96 well-plates

Aliquots of 140 µL of the freshly prepared mono- and co-culture cell suspensions were distributed in the wells of a 96 well-plate (Cyto One, tissue culture treated, Catalog No: CC7682-7596), in six to seven technical replicates. Six to seven wells with the same sterile medium were incubated as controls for sterility. The plate was then incubated at 30°C in a plate reader (BioTek, plate reader, Synergy H1), for up to 48 h. Plates were continuously shaken (double orbital, 282 cpm, slow orbital speed). Absorbance (OD_600_) and fluorescence (GFP - 480/510 Ex/Em, and mCherry – 580/610 Ex/Em) were measured every 30 min in each cultivation well. After the incubation, the plates were placed on ice and sampled for flow-cytometry counting (see below).

### Microfluidic encapsulation of cells in droplets and culture procedure

Similarly, precultured mono- and co-culture cell suspensions were used for microfluidic encapsulation (Fig. 1), but now with an estimated 5×10^7^ cells per mL, in order to obtain 1–3 founder cells at start within a 35 pL-volume. An aliquot of 500 µL of diluted cell suspension (mono- or co-culture) was taken up in a 1-mL syringe (Omnifix 1 mL, U-100 Insulin), and 1 mL of HFE 7500 Novec fluorinated oil containing 2 % (w/w) of fluorosurfactant (RAN Biotechnologies, Inc.) was loaded in another one. Oil-dissolved surfactant stabilizes formed aqueous droplets and prevents them from coalescing. Syringes with the aqueous cell suspensions and with the oil were mounted on two separate syringe pumps (Harvard Apparatus, Pump 11 Elite / Pico Plus Elite OEM Syringe Pump Modules) to inject the liquids into a droplet maker microfluidic chip ^42^ at flow rates of 8 µL min^−1^ and 20 µL min^−1^, respectively. The droplet maker chip (custom-produced by Wunderlichip GmbH, CH-8037 Zürich, Switzerland) used has a 40×40×40 µm junction (see Fig. S1), generating monodispersed droplets with a diameter of 40 µm. Formed droplets were collected for 10 min in a 1.5 mL Eppendorf tube, corresponding to a total volume of 80 µL of droplets. The Eppendorf tube was prefilled with 250 µL phosphate-buffered saline (PBS) to prevent the droplets from evaporating. Eppendorf tubes with the droplets were kept on ice until being incubated at 30°C to maintain the starting cell concentrations during the collection of the droplets from the different conditions tested in parallel (mono- and co-cultures). After a first timepoint imaging (t = 0 h), droplets were incubated at 30°C and sampled at different intervals (Table S4). After the final incubation time, the vials with the droplets were placed on ice before coalescing all droplets and counting cell numbers by flow cytometry (see below).

### Droplet sampling

Droplet cultures were sampled at the start of the incubation, and after 17, 24 or 48 h (depending on the condition, Table S4). An aliquot of 1.5 µL was retrieved from the droplet emulsion, which was transferred by micro-pipette into a 5–µL HFE 7500 oil layer inside a chamber observation slide (Countess chamber slide, Invitrogen C10228). Then, another volume of 5 µL of oil was added to the chamber to disperse droplets in a monolayer. Droplets were imaged at 3-5 random individual positions with a Leica DMi4000 inverted epifluorescence microscope (*P. putida - P. veronii* experiments) or a Nikon Ti2000 inverted epifluorescence microscope (for the four other strains experiments), a Flash4 Hamamatsu camera, a 20x objective (Leica, HI PLAN I 20x/0,30 PH1, with *P. putida-P. veronii*, or a Nikon CFI S Plan Fluor ELWD 20XC MRH08230, with the four other strains), in Bright field (exposure time = 5 ms), red (exposure time = 400 ms) and green fluorescence (exposure time = 600 ms, for *P. putida* and 400 ms for the other strains). Images were collected as 16-bit .TIF files and further analyzed with a custom-made Matlab script to segment droplets and cells in droplets.

### Timelapse imaging of cell growth in droplets

In select droplet experiments (Table S4), we followed the growth of cells in individual droplets over time by timelapse microscopy in an observation chip (Fig. 6A, chip design adopted from ref ^50^, custom-produced by Wunderlichip GmbH). This polydimethylsiloxane (PDMS) print was directly mounted on a 1-well chambered Coverglass (Nunc™ Lab-Tek™ II Chambered Coverglass, Thermo Fisher, Cat number: 155360PK), to be able to immerse the chip during the observation. Before loading, the “chambered” chips were placed in a vacuum chamber for 20 min to extract any gas contained in the PDMS, thus preventing the appearance of air bubbles during the incubation (we acknowledge that this potentially reduces the level of available oxygen to the cells). The chip was then filled and immersed in deionized filtered (0.22–µm) water overnight. One hour before loading the droplets, the chip flow lines were emptied from the water and refilled with HFE 7500 oil, and the immersion chamber of the chip was filled with 1 mL of HFE 7500 oil, on top of which was placed 4.5 mL of deionized water, to limit oil and droplet evaporation during the incubation and imaging. Cell suspensions were encapsulated into droplets following the same procedure as explained above, but now the production chip outlet was directly connected by teflon (PTFE) tubing to the inlet of the (immersed) observation chip. An aliquot of 20 µL of HFE 7500 oil was pipetted inside the observation chip inlet (using P20 tips) to allow good separation of the incoming droplets (this was done under live observation of the observation chip with an inverted microscope, to verify droplet separation). Droplets accidentally leaking into the chamber were removed by pipetting. Finally, aliquots of 30 µL of HFE 7500 oil were pipetted into the two outlets of the observation chip, to prevent water from entering the chip during incubation and imaging. The immersion chamber was then closed and sealed with parafilm. The height of the chamber in the observation chip is 10 µm, which causes droplets to squeeze, and to almost completely fall within the focal depth range of the 20× objective. The chip was mounted on a Nikon Ti2000 inverted epifluorescence microscope with a programmable stage and was imaged every 10 min in the three channels (BF, GFP and mCherry) as before, at the same individual positions set with the imaging control (Micromanager software 1.4.23). Images were exported as 16-bit .TIF files.

### Image analysis

TIF-images were processed in a custom-made Matlab script (see Data availability), which segments all droplets per image and all fluorescent objects per droplet. The script then calculates the sum of all fluorescent objects per droplet (in pixel area), which is multiplied by their mean fluorescence intensity, to obtain a ‘total fluorescent signal’ (AF, see Fig. 1). We use the AF-value per droplet as a proxy for the biomass production of the strain identified by its specific fluorescence (Table S3), under the assumption that the more cells there are in a droplet, the higher their fluorescent signal will be (Fig. 4). Depending on the fluorescence intensity displayed by the cell-strain-pairs, the raw fluorescent signals were log_10_-, square-root or median-transformed for display of potential subpopulations. The distributions of fluorescent signals are then analysed across all droplets, and across independent biological replicates. For timelapse experiments, images were segmented and processed in the same way as above, with the difference that a customized rolling ball algorithm was applied to compensate in the image segmentation for fluorescence variations existing among cells and bleaching of the signal along the timelapse. Additionally, the droplets were tracked between time frames, by comparing the distances of the centroid for every droplet between frame *t* and the next frame *t+1*. Droplets with minimum centroid distances were assumed to be the same on frame *t+1* as on frame *t*. Tracking of individual droplets was then manually controlled and corrected with the help of generated movies displaying the tracking ID attributed to each droplet over time. In this way, biomass development can be plotted per droplet over time, and the variation among droplets can be quantified.

### Fusing droplet emulsions for flow cytometry cell counting

Droplets from a single Eppendorf emulsion experiment were fused to produce a single aqueous phase, in which the total cell yield could be counted by flow cytometry. First, the extra HFE oil that settled below the droplet emulsion was removed by pipetting. To the remaining PBS and droplet emulsion layer, an approximate equivalent volume was added of HFE oil containing 1H,1H,2H2H-perfluoro-1-octanol (5 g solution Sigma-Aldrich, further diluted 4 times in HFE 7500 oil). This breaks the emulsion and fuses the droplets into a single aqueous phase. The resulting droplet-cell-PBS aqueous phase was transferred into a new Eppendorf vial and its volume was measured from the micro-pipette directly.

### Flow cytometry counting of cell population sizes

Cell numbers in liquid suspensions from fused droplet emulsions or mixed liquid suspended cultures in 96-well plates, or precultures, were quantified by flow cytometry. Liquid cell suspensions were tenfold serially diluted in PBS (down to 10^-3^) and fixed by adding NaN_3_ solution to a final concentration of 4 g L^−1^ and incubating for max 1 day at 4 °C until flow-cytometry processing. Volumes of 20 µL of fixed samples were aspired in a Novocyte flow-cytometer (Bucher Biotec, ACEA biosciences Inc.) at 14 µL min^−1^. Events were collected above general thresholds of FSC = 500 and SSC = 150 to distinguish cells from particle noise, and gates were defined to selectively identify the strains from their fluorescent markers (Table S3, see gating example in Fig. S10). The Novocyte gives direct volumetric counts, which were corrected for the dilution. To convert cell counts from droplet suspensions to equivalent cell concentrations per mL, we considered the proportion of empty droplets from imaging and the extra volume of 250 µL of PBS before droplet collection, as follows:

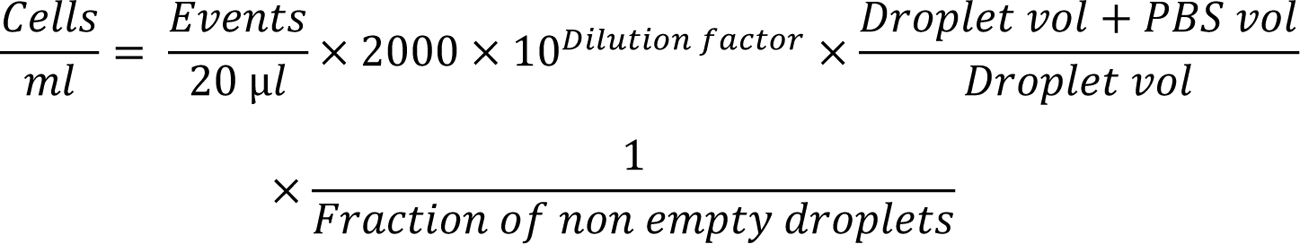

The multiplication by 2000 includes the 2-fold dilution when fixing the sample with NaN_3_ solution, and the conversion to a per-mL concentration.

### Calculation of maximum specific growth rates, lag times and time to first population doubling

Average growth rates and lag times of strains in suspended liquid culture were inferred from the ln-transformed strain-specific fluorescence increase in mono-cultures grown in 21C MM with their specific carbon substrate (as described above), each in 6-7 replicates. To average, we calculate the slope over a sliding window of five consecutive timepoints during the first 10 h, retained only slopes with a regression coefficient > 0.97; and reported the mean of those slopes as the maximum specific growth rate. Lag times were fitted from the complete (fluorescence) growth curve using a logistic function, and converted to *time to first population doubling* as the sum of the lag time (in h) plus the inverse of the logarithmic fitting constant multiplied by ln(2). (In absence of lag time the time to first population doubling is the inverse of the maximum specific Monod growth rate.)

To calculate growth rates from fluorescence in single droplets, we deployed a manual interactive plot of ln-transformed values of the summed fluorescence signal (the product of segmented area and the average strain-specific fluorescence in that area) over time, identifying the start and ends of the ln-linear range, and the *lag* time being the time between the start of the imaging series and the start of the ln-linear range. The maximum specific growth rate in the droplet was then taken as the slope over the entire identified ln-linear range. Since we did not segment individual cells, the summed fluorescence signal per droplet is a proxy for their ‘biomass’, and we report a Monod-type maximum specific growth rate.

### Mathematical model for population growth in droplets

We adapted a previously developed mathematical framework ^49^ to simulate the growth of *P. putida* and *P. veronii* populations in 35 pL droplets with nutrients (10 mM succinate). The initial resource concentration (R_0_) is homogeneously distributed among all droplets and can not diffuse between droplets. The chemical reactions inside each droplet are similar to the bulk population model in Ref. ^49^, however, each founder cell follows its own differential growth equation, and includes possible kinetic variation. Growth of each founder cell *i* in droplet *j* thus follows the general reaction,

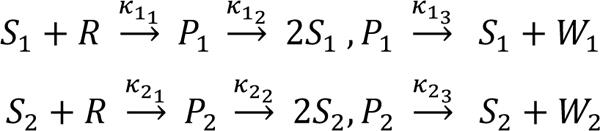

where *S* is the bacterial species, *R* represents the resource, *P* is the cell-resource intermediate state and *W* any non-used metabolic side products. For simplicity, we did not consider cross-feeding effects. Each founder cell *i* has its own lag time 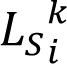 where *i* ∈ 1, 2 is the species index and *k* ∈ ℕ the founder cell index. Therefore, the differential equations for species *S_1_* or *S_2_* present in the droplet j are,

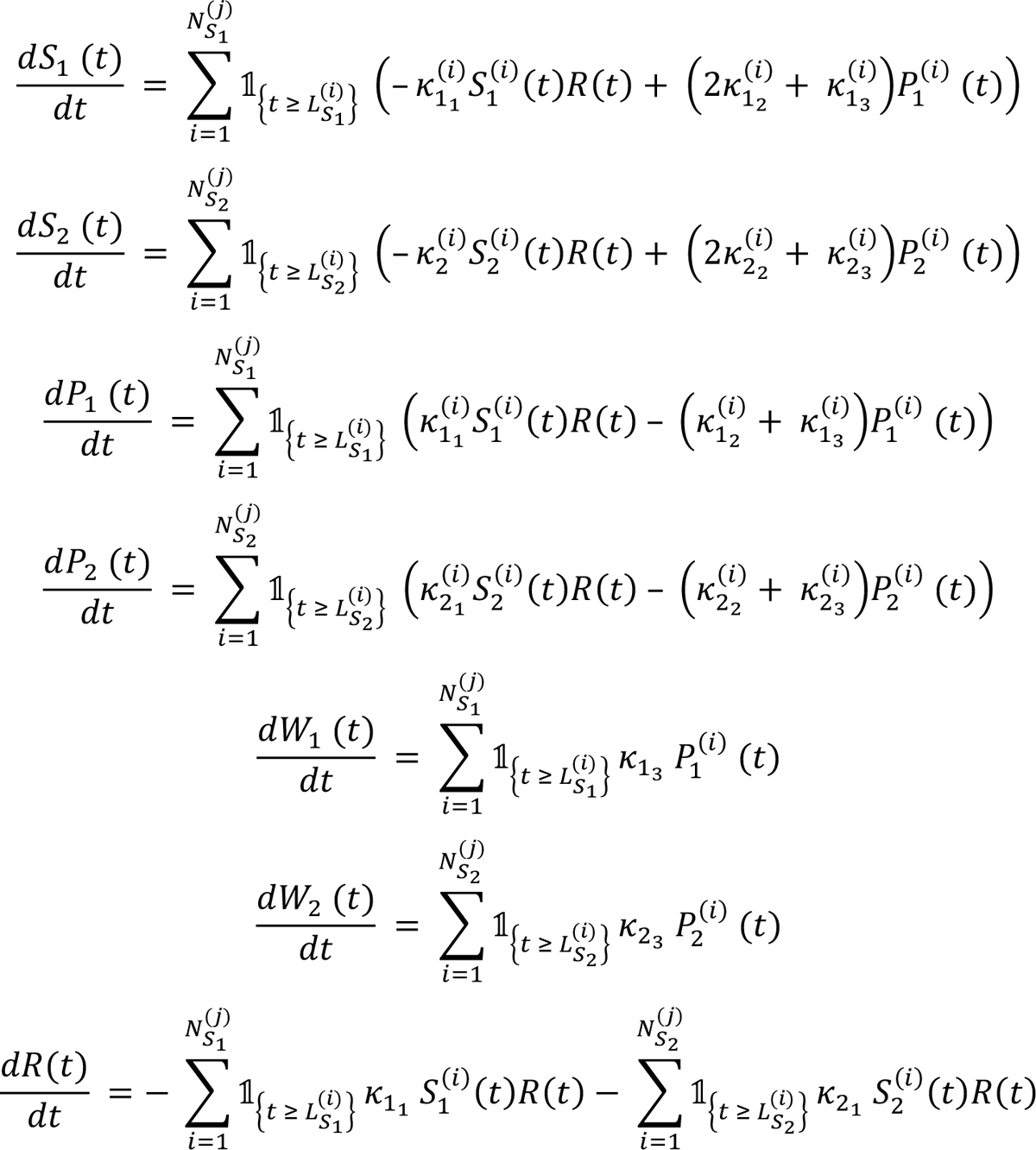

where 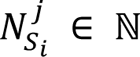 is the number of initial cells of species *i* ∈ 1,2 in the droplet *j*.

Cells of both species are Poisson-distributed across the droplets with an average of 3 cells per species. Individual growth rates and lag time parameters were sampled from a generated Gamma-distribution of *P. putida* and *P. veronii* growth parameters, inferred from mono-culture OD_600_ curves with an Monte-Carlo Metropolis Hasting algorithm centred on the mean (Fig. S3, analysis as described in reference ^49^). Cell-to-cell growth kinetic variance was taken from the timelapse growth curve measurements in droplets with single-founder cells of *P. putida* and/or *P. veronii* (Fig. S11). We also included a 15% chance for a cell to have a *null*-growth rate, to account for growth-impaired cells that we observed from droplet imaging (Fig. 2E). Varying the heterogeneity in growth properties in Figures 7F-H among founder cells thus consisted in increasing or decreasing the initial variance of the parameter gamma distributions.

### Statistical analysis and reproducibility

All experiments were carried out in biological triplicates (quadruplicate incubations for the substrate independence scenario). For each biological replicate, liquid-suspended cultures comprised 6-7 cultivation wells as technical replicates. Biological replicates of fragmented droplet cultures comprised one separate emulsion incubation each, except in one of the replicates of the substrate competition experiment, for which a triplicate emulsion was generated, to assess and show the technical reproducibility of droplet cultivation experiments (Fig. S12). Each emulsion sample was then imaged at 5-20 positions (technical replicates), to obtain 100-1000 droplets per mono- or co-culture and treatment. Flow cytometry counts (Fig. 2C & D, Fig. 3C, Fig. 4C) show the means of all technical replicates within each biological replicate. Each suspension from a cultivation well or fused droplet emulsion was counted three times by the flow cytometer, from which the mean was taken. T-tests were conducted to compare mean cell-counts in flow cytometry. Normality in the data was verified with a Shapiro-Wilk’s test, and variance homogeneity was verified with a Fisher test. Median, top-10 or low-10 percentile productivities of each species in mono vs mix droplets were compared using Wilcoxon rank-sum or sign-rank tests (when taken across multiple time points). Depending on the data, we tested against a *null* hypothesis of sample means or ranked values being indifferent, or being higher or lower (i.e., a left or right tail). Tests were implemented in *R* version (within *Rstudio* version 2022.07.01) or in MATLAB (MathWorks, Inc. version R2021b).

To deduce strain interactions, we compare observed mixed droplet growth with the expected mixed growth from a *null* model based on probability distributions generated from the corresponding mono-culture droplet growth (*i.e.*, assuming no interactions). The model uses the probability distributions for productivities of each of the strains in pairs at each sampled time point, simulated five times for the same number of pairs as the number of observed droplets. Expected and observed paired droplets are then counted in a productivity grid (*e.g.*, as in Fig. 3F), and summed fractions across relevant grid regions (*e.g.*, higher than 1.5 times the median) are compared across replicates (typically, three biological replicates and five simulation replicates). P-values are then derived in an *ANOVA* comparison including all fractions, followed by a *post-hoc* multiple test (Fig. 3F), or by Wilcoxon sign-rank test in case of comparing multiple time points (e.g., Fig. 4F), as implemented in MATLAB.

To estimate the proportion of mixed droplets in which Pf-5 might have been killed (lysed) by CHA0 tailocins, we deployed variations in the specific median fluorescence background originating from Pf-5. We first calculate the standard variation in Pf-5 background fluorescence from Pf-5 ‘solo’ droplets, corrected for the Pf-5 biomass (*i.e.*, segmented area), which is multiplied by 2.5 as a boundary for the outlier range. This outlier range definition was then imposed on the Pf-5 specific fluorescence in mix droplets with CHA0 (and in CHA0 droplets where no Pf-5 area can be distinguished, assuming they may all be lysed). Outlier fractions were corrected for the total observed droplets, and compared to the outlier fractions observed for Pf-5 solo droplets (*i.e.*, as in Fig. 5E and 5F), using one- or two-tailed two-sample *t-*testing of replicate values.

The effect of founder cell census on the variance of growth kinetic parameters in strain-paired droplets was examined using a generalized linear mixed effect model (*glme*, as implemented in MATLAB 2021a), using measured maximum specific growth rates, lag times and starting cell ratios as variables. Droplets with quasi-null growth rates (which were also characterized by a lag time above 20 h) were removed for the analysis. The relationship of individual growth rate and lag time variance as a function of founder cells was further explored using a Brown-Forsythe test implemented in *R*, which tests the homogeneity of variances between groups without assuming the normality of the data.

## Data availability

All processed droplet productivity data, numerical data values underlying figure elements and custom R/MATLAB scripts used for image analysis will be made available from a single downloadable link on Zenodo.

## Acknowledgements

The authors thank Yolanda Schaerli and Florian Baier for their initial help in the droplet microchip setup, Vladimir Sentchilo for the genetic tagging of *Pseudomonas* strain L15, Bouke Bentvelsen and Guillaume Lieb for helping with image segmentation and Martin Ackermann for helpful discussions.

