## Supplementary material for "Fragmented micro-growth habitats present opportunities for alternative competitive outcomes": Batsch_supplementaryinformation.pdf

**Supplementary tables**

Table S1 - Strain-specific fluorescence distributions of solo- or mix droplets with *S. wittichii* RW1 and *Pseudomonas* sp. Leaf15 under substrate indifference conditions

Table S2: Generalized linear mixed effects analysis for paired productivities of *P. putida* and *P. veronii* in timelapse imaged droplets

Table S3: Strains and pre-culture procedures

Table S4: Strains and media conditions for each experiment

**Supplementary figures**

Figure S1: Encapsulation of cells in picoliter droplets by microfluidic operation.

Figure S2: Distribution of *P. putida* and *P. veronii* cells after encapsulation in pL droplets.

Figure S3: Biomass growth in suspended monocultures of *P. putida* and *P. veronii*.

Figure S4: Fluorescence growth curves of *P. putida* and *P. veronii* cultivated on D-mannitol and putrescine.

Figure S5: Substrate preferences of *Pseudomonas* L15 and *Sphingomonas wittichii* RW1.

Figure S6: Fragmented productivities of *Pseudomonas* Leaf15 (L15) and *Sphingomonas wittichii* RW1 in pL droplets over time.

Figure S7: Stationary productivity of *P. putida* and *P. veronii* in droplets vs lag time.

Figure S8: Paired growth trajectories of *P. putida* and *P. veronii* in time-lapse imaged droplets.

Figure S9: Influence of founder cell census on droplet population growth kinetics.

Figure S10: Gating strategy of events detected in flow cytometry.

Figure S11: Distributions of growth kinetics parameters measured from droplets with a single founder cell of *P. putida* or *P. veronii* imaged on a timelapse.

Figure S12: Technical reproducibility of droplet cultivation procedures.

**Supplementary movies**

Movie S1: Focal movie on one droplet with simultaneous growth of both *P. protegens* CHA0 and Pf-5 with 10 mM succinate.

Movie S2: Focal movie on one droplet with lysis of *P. protegens* Pf-5 cells during co-culturing with CHA0 on 10 mM succinate.

**Supplementary references**

**Supplementary tables:**

**Table S1: Strain-specific fluorescence distributions of solo- or mix droplets with *S. wittichii* RW1 and *Pseudomonas* sp. Leaf15 under substrate indifference conditions.**

| Comparison | *S. wittichii* RW1 | | *P*-value[1] | *Pseudomonas* sp. Leaf15 | | *P*-value |
|---|---|---|---|---|---|---|
| | Solo droplets | Mix droplets | | Solo droplets | Mix droplets | |
| Mean of replicate medians[2] | 0.8564 | 0.9079 | 0.0488 | 0.9306 | 0.9979 | 0.0019 |
| Mean of replicate 90$^{th}$ percentile | 1.8395 | 1.9058 | 0.1797 | 1.4989 | 1.6074 | 0.0019 |
| Mean of replicate 10th percentile | 0.3488 | 0.3332 | 0.2129 | 0.5437 | 0.5947 | 0.0273 |
| Mean fraction of top 10 percentiles | 0.0995 | 0.1158 | 0.125 | 0.1000 | 0.1325 | 0.0019 |
| Mean fraction of low 10 percentiles | 0.0995 | 0.1213 | 0.0820 | 0.1000 | 0.0837 | 0.0488 |

1) Sign-rank test implemented in MATLAB

2) Three biological replicates from three sampling time points combined

**Table S2: Generalized linear mixed effects analysis for paired productivities of *P. putida* and *P. veronii* in timelapse imaged droplets**.

| **Fixed effects coefficients** | **Estimate** | **p-value** | **SE** | **tStat** | **DF** |
|---|---|---|---|---|---|
| (Intercept) | -0.96152 | 0.52747 | 1.5165 | -0.63405 | 102 |
| PVE growth rate | -21.833 | $9.4472 \times 10^{-18}$ | 2.0947 | -10.423 | 102 |
| PVE lag time | 1.4238 | $3.6464 \times 10^{-33}$ | 0.079734 | 17.857 | 102 |
| PPU growth rate | -0.39277 | 0.91429 | 3.6405 | -0.10789 | 102 |
| PPU lag time | -0.23682 | 0.031472 | 0.10858 | -2.1811 | 102 |
| Starting cells ratio | 0.58324 | 0.011955 | 0.22789 | 2.5593 | 102 |

The contribution of population kinetics and starting cell ratios to the stationary productivity ratio of *P. putida* (PPU) and *P. veronii* (PVE) in droplets (10 mM succinate) was assessed via a

GLME analysis. SE gives the Standard Error for the estimated value (Estimate) of each Fixed effects coefficient. DF gives the Degree of Freedom corresponding to the t-statistic (tStat).

**Table S3: Strains and preculturing procedures**

| **Strain name** | **Agar plate media** | **Pre-culture media** | **Pre-culture time** | **Final $OD_{600}$ (before mixing)** | **Fluorescent marker for identification** | **Source or reference** | **Gate in flow cytometry** |
|---|---|---|---|---|---|---|---|
| *P. putida* (PPU) | NA | 21C MM + 10 mM succinate | 16h | 0.02 | eGFP | Carraro et al. 2020 (1) | FSC-H, FITC-H |
| *P. veronii* (PVE) | NA | 21C MM + 10 mM succinate | 16h | 0.05 | mCherry | Dubey et al. 2021 (2) | FSC-H, PE-TexasRed-H |
| *Pseudomonas sp.* Leaf15 (L15) | R2A | 21C MM + 10 mM succinate | 48h | 0.02 | mScarlet-I | Helfrich et al. 2018 (3), This study | FSC-H, PE-TexasRed-H |
| *Sphingomonas wittichii* (RW1) | R2A | 21C MM + 4 mM salicylate | 48h | 0.02 | eGFP | Coronado et al. 2015 (4) | FSC-H, FITC-H |
| *P. protegens* CHA0 (CHA0) | NA | 21C MM + 10 mM succinate | 16h | 0.05 | GFP2 | Vacheron et al. 2021 (5) | FSC-H, FITC-H |
| *P. protegens* Pf-5 (Pf-5) | NA | 21C MM + 10 mM succinate | 16h | 0.05 | mScarlet-I | This study | FSC-H, PE-TexasRed-H |

**Table S4: Strains and media conditions for each experiment**

| **Experiment scenario** | **Strains involved** | **Cultivation media** | **Droplet sampling times** | **Droplet time-lapse** |
|---|---|---|---|---|
| Substrate competition | PPU; PVE | MM C21 + 10 mM succinate | 0, 24 h | yes |
| Substrate independence | PPU; PVE | MM C21 + 10 mM d-mannitol + 6.67 mM putrescine | 0, 24, 48 h | no |

| | | | | |
|---|---|---|---|---|
| Substrate independence + growth inhibition | L15; RW1 | MM C21 + 4 mM succinate + 1.5 mM salicylate | 0, 17, 24, 48 h | no |
| Substrate competition + tailocin interaction | CHA0; Pf-5 | MM C21 + 4 mM succinate | 0, 24, 48 h | yes |

**Supplementary figures:**

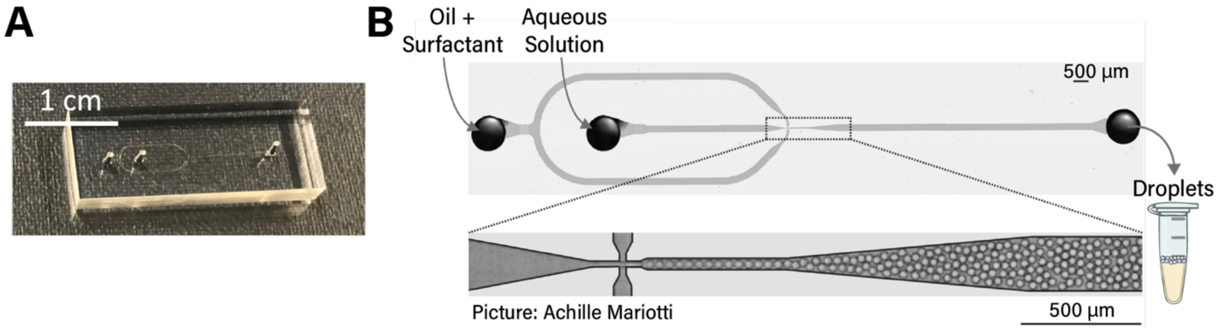


**Figure S1: Encapsulation of cells in picoliter droplets by microfluidic operation**. A) Chip design from Duarte *et al. (2017)*. B) The junction between oil and aqueous flows has a 40 µm x 40 µm x 40 µm dimension, allowing for the generation of monodispersed 40 µm-sized droplets. Droplets are directly collected in an Eppendorf tube. Pictures shared with the courtesy of Achille Mariotti.

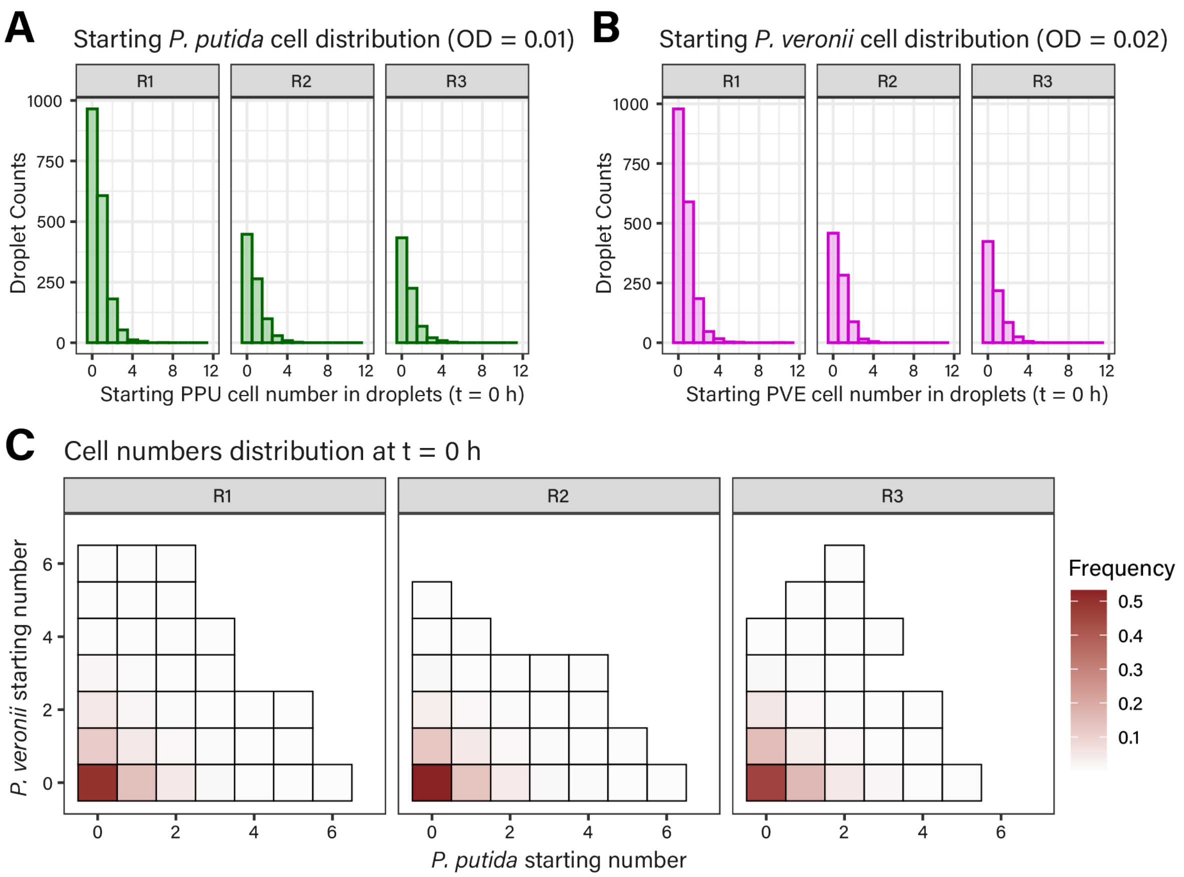


**Figure S2: Distribution of *P. putida* and *P. veronii* cells after encapsulation in pL droplets.** A) Distribution for *P. putida* (PPU) and B) *P. veronii* (PVE) cells at the beginning of the substrate competition experiment in the mixed culture droplets (t = 0 h). C) Heatmap of paired frequency distributions of *P. putida* and *P. veronii* cells in droplets (t = 0 h).

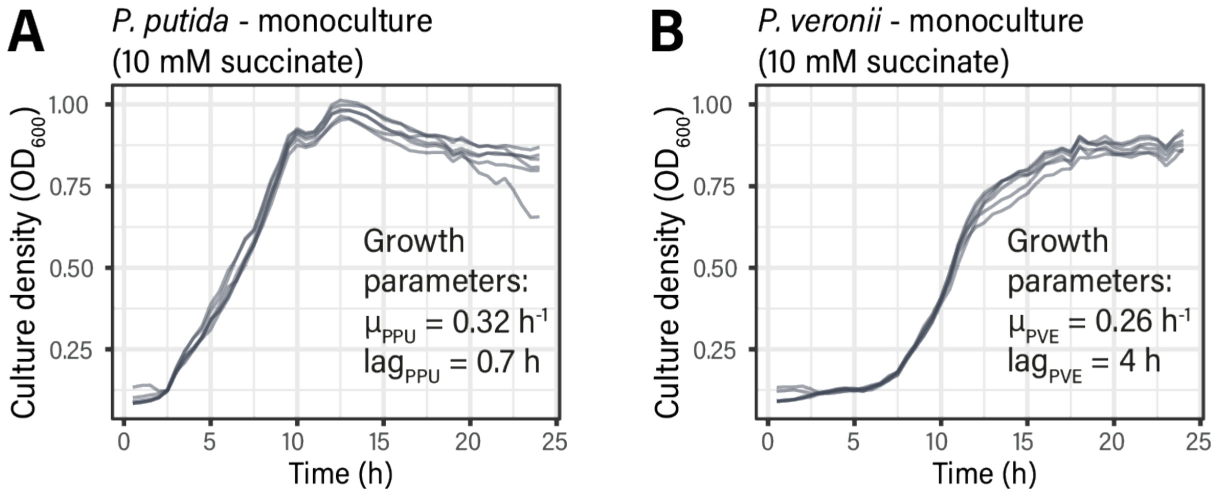


**Figure S3**: **Biomass evolution in suspended monocultures of *P. putida* and *P. veronii***. Growth curves of *P. putida* (A) and *P. veronii* (B) monocultures on 10 mM succinate was used to predict inherent growth kinetic parameters of both species on succinate with an MCMC Metropolis Hasting algorithm method.

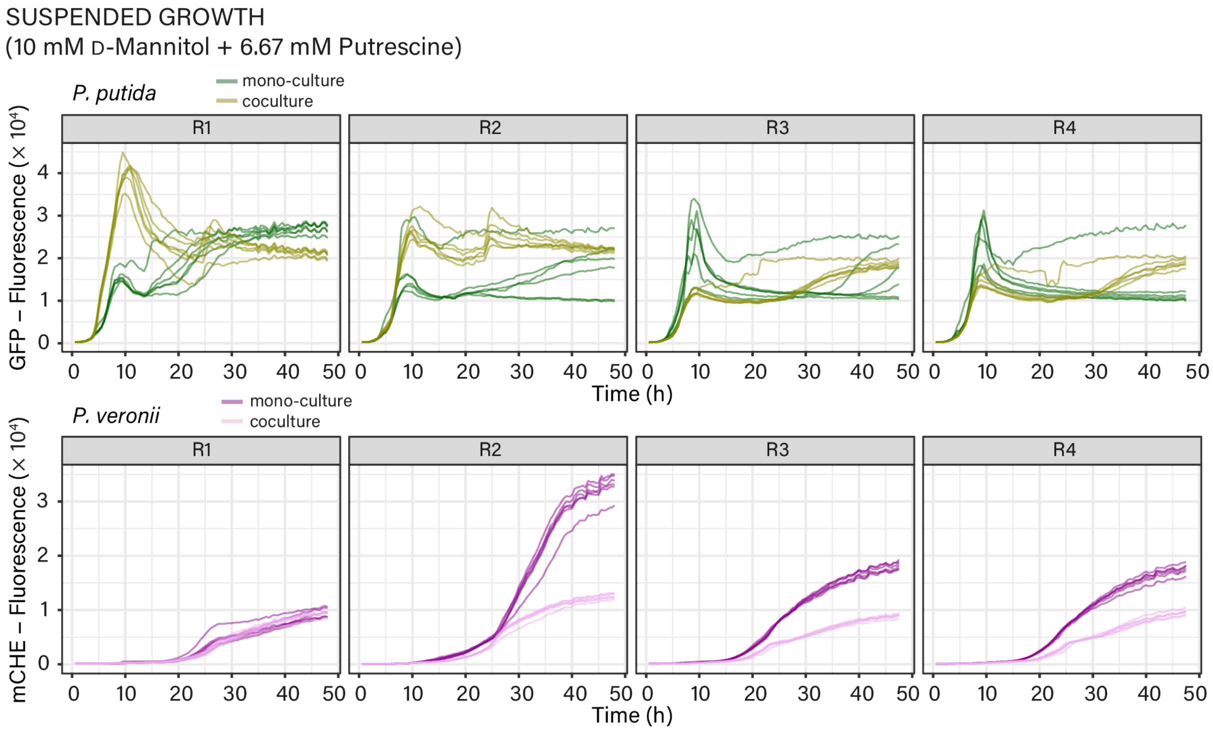


**Figure S4: Fluorescence growth curves of *P. putida* and *P. veronii* cultivated on D-mannitol and putrescine**. Both strains were grown together or separately in 96-well plates. Each panel corresponds to one biological replicate (with n = 6 technical replicates for each).

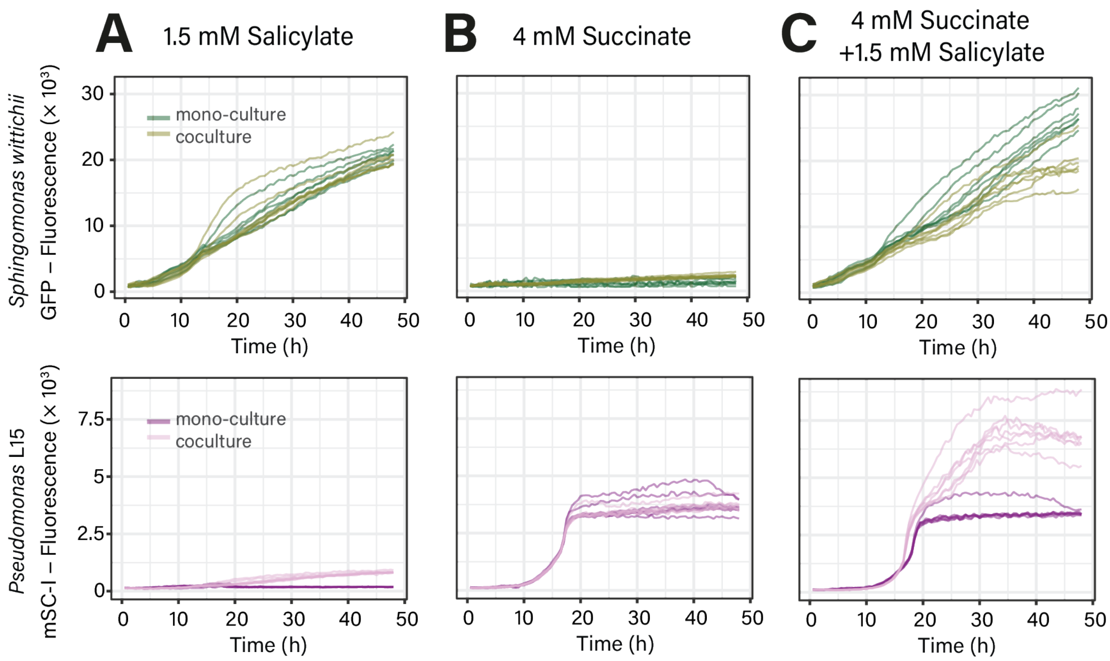


**Figure S5: Substrate preferences of *Pseudomonas* L15 and *Sphingomonas wittichii* RW1**. Fluorescence growth curves of the two strains grown together or separately in 96-well plates on A) salicylate, B) succinate, or C) both salicylate and succinate, show the independent usage of salicylate by RW1 and of succinate by L15 , with a potential cross-feeding from RW1 to L15 (n = 7 technical replicates).

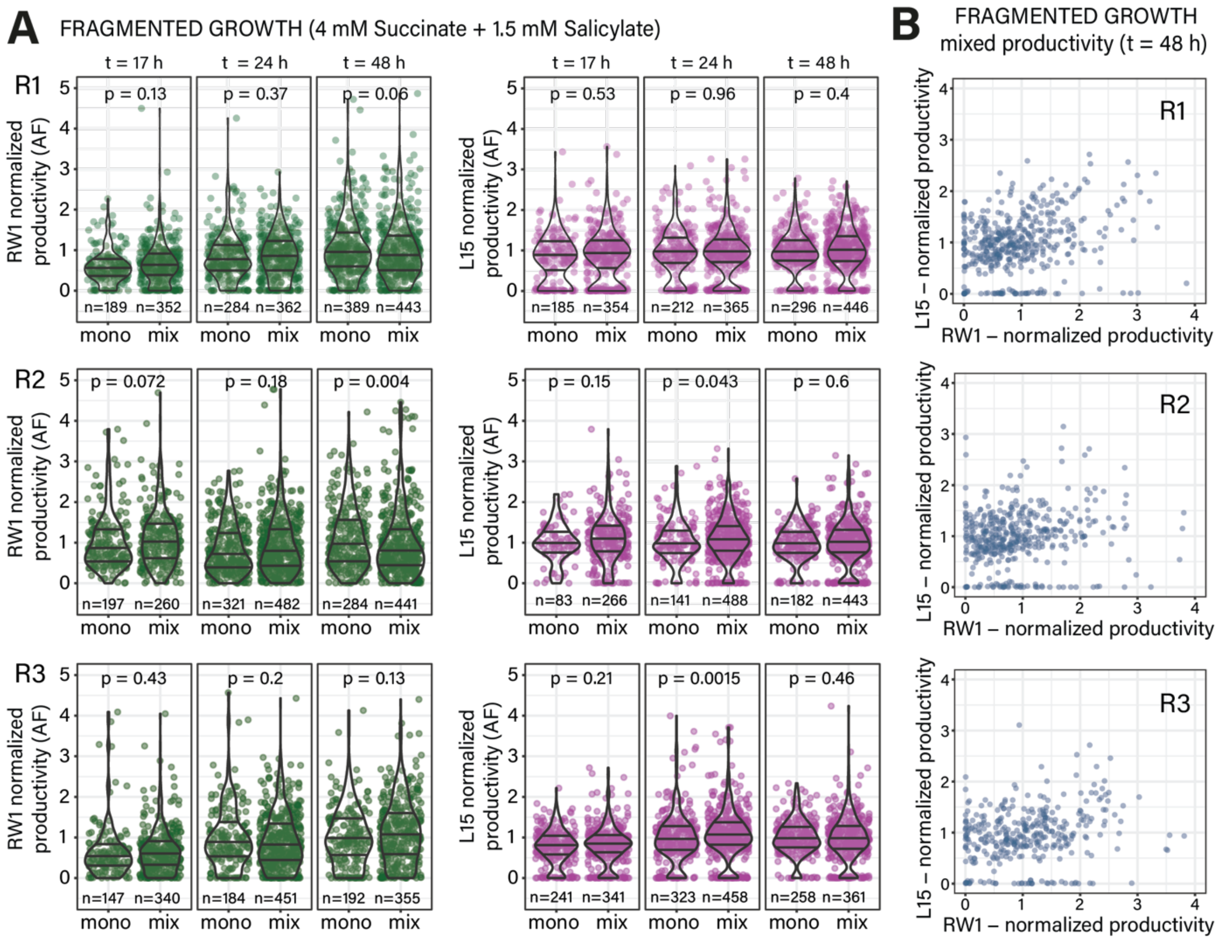


**Figure S6: Fragmented productivities of *Pseudomonas* Leaf15 (L15) and *Sphingomonas wittichii* RW1 in pL droplets over time.** A) Productivities of the two strains in solo and mixed droplets show generally non-significant differences between the two conditions over time. Droplets counts is indicated under each jitter-plot. B) Paired productivities of L15 and RW1 in mixed droplets after 48 h. R1, R2 and R3 denote the three biological replicates of the experiment. P-values from two-sided Wilcoxon rank sum tests.

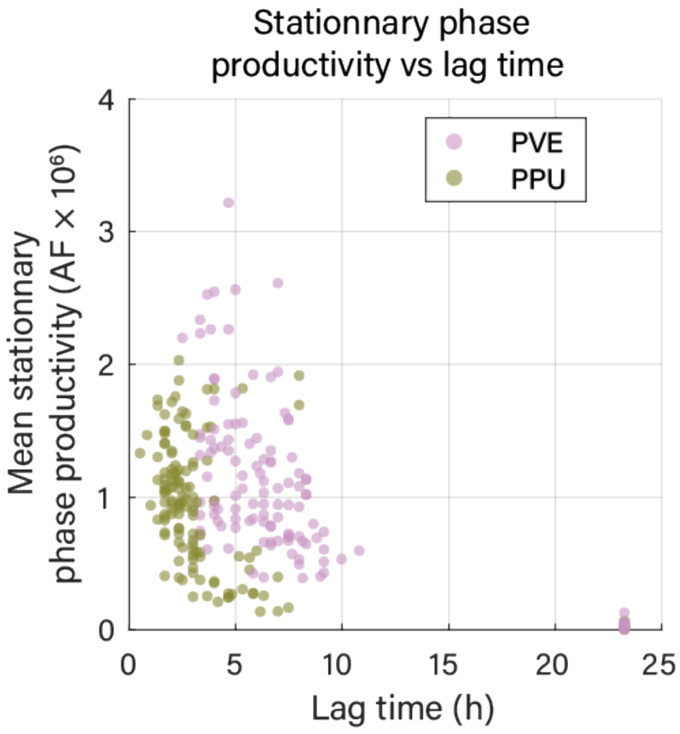


**Figure S7: Stationary productivity of *P. putida* and *P. veronii* in droplets vs lag time**. Data points from mixed droplets of *P. putida* (PPU) and *P. veronii* (PVE) growing in competition for succinate during timelapse imaging experiment.

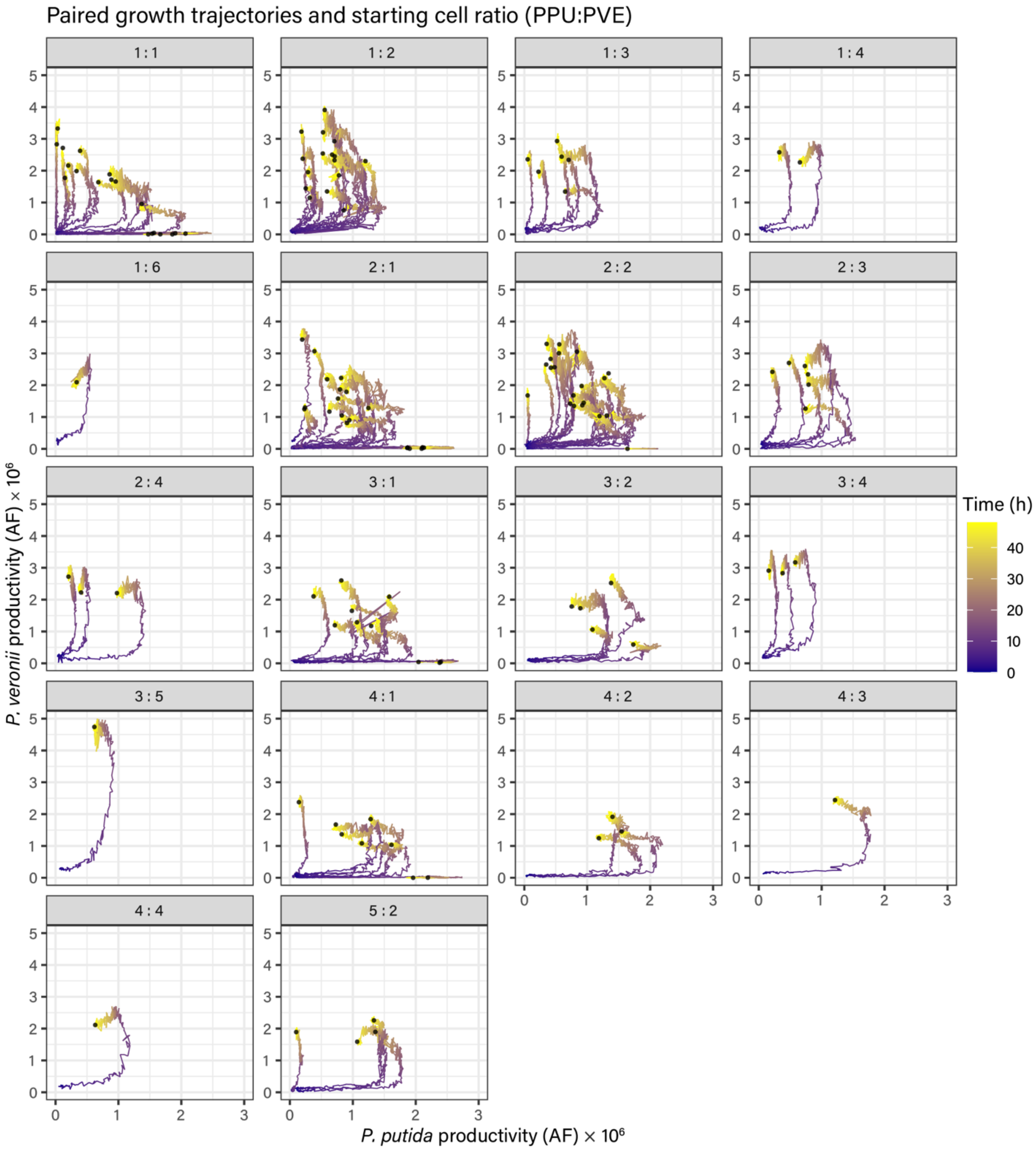


**Figure S8: Paired growth trajectories of *P. putida* and *P. veronii* in time-lapse imaged droplets.** Each facet indicates the starting number of *P. putida* (PPU) and *P. veronii* (PVE) cells (PPU:PVE) and thus regroups paired growth curves of droplets starting in the same conditions.

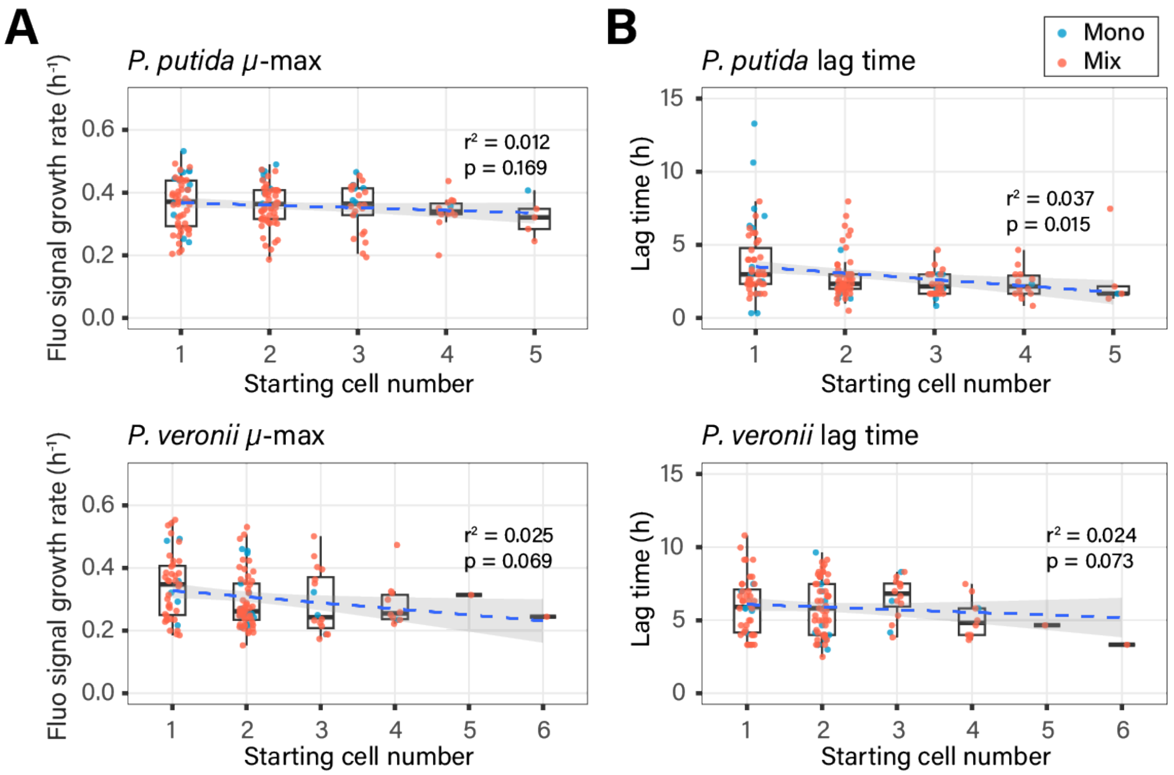


**Figure S9: Influence of founder cell census on droplet population growth kinetics.** A) measured growth rates and B) lag times for *P. putida* and *P. veronii*, measured in droplets from the AF-fluorescent signal evolution in droplets, as a function of each species' founder cell census. No significant change with the increasing number of founder cells is detected (except an average decrease for the lag times of *P. putida*).

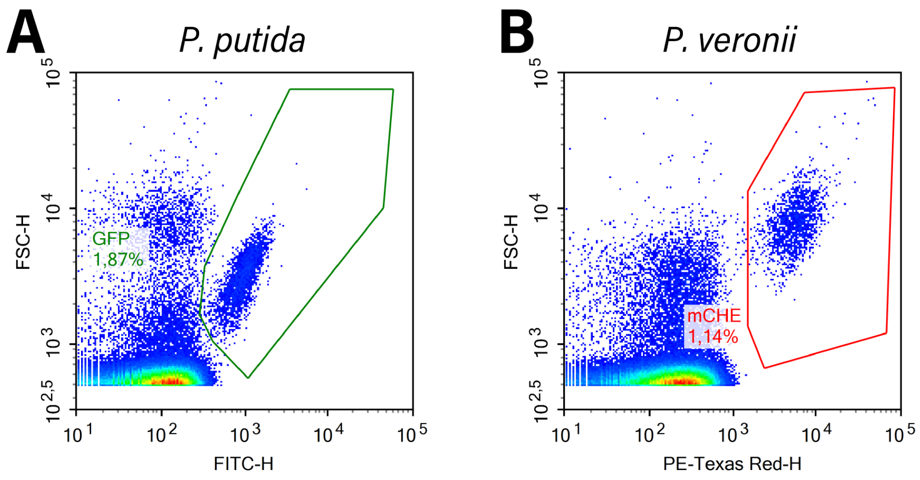


**Figure S10: Gating strategy of events detected in flow-cytometry**. Examples for *P. putida* (A) and *P. veronii* (B)

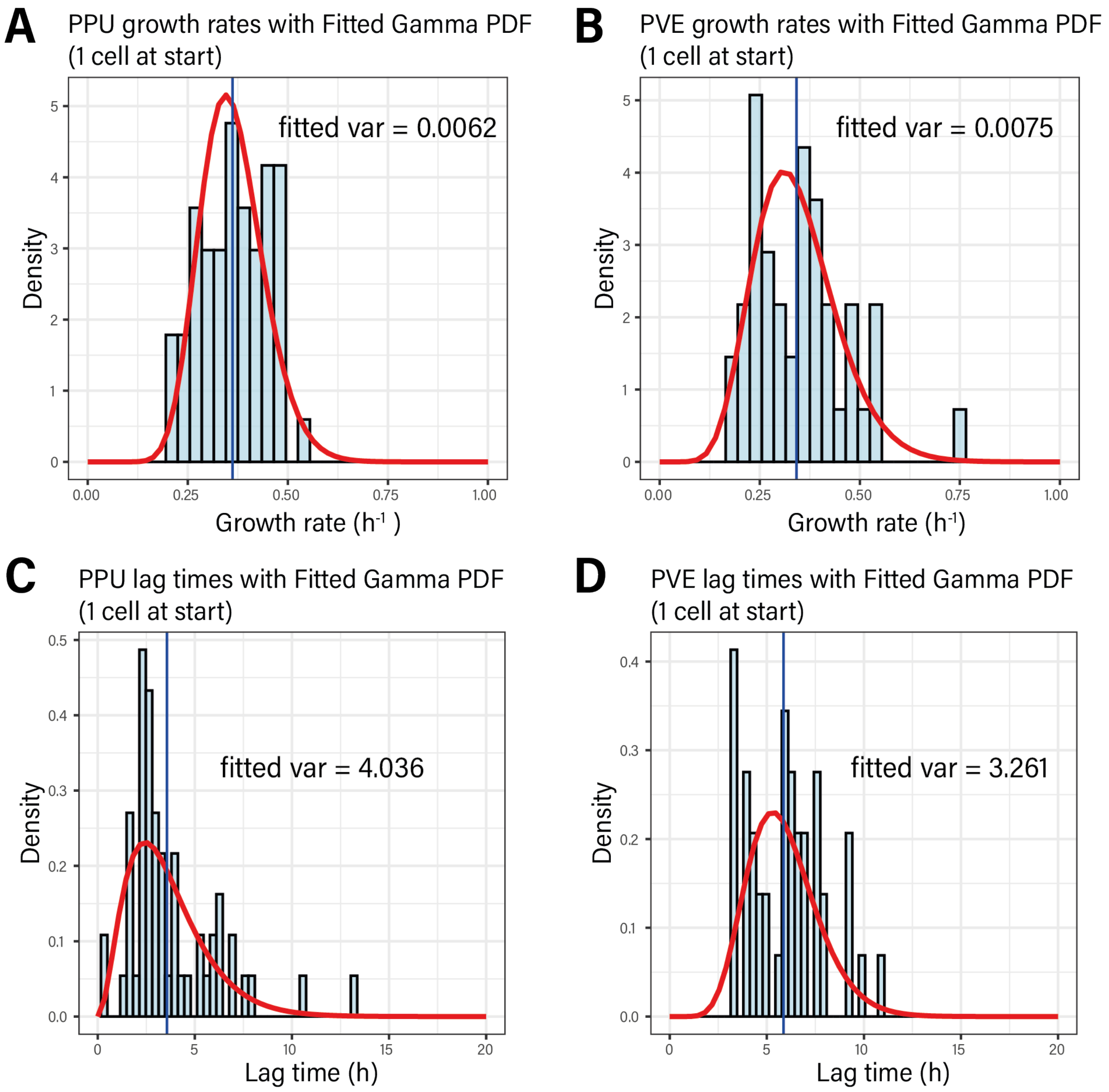


**Figure S11: Distributions of growth kinetics parameters measured from droplets with a single founder cell of *P. putida* or *P. veronii* imaged on a timelapse.** A) Growth rates of *P. putida* (PPU) and B) *P. veronii* (PVE). C) Lag time of *P. putida* and D) *P. veronii*. Kinetic parameters displayed here combine mono and mixed droplets. Displayed variances were inferred by fitting Gamma distributions to the parameter distributions.

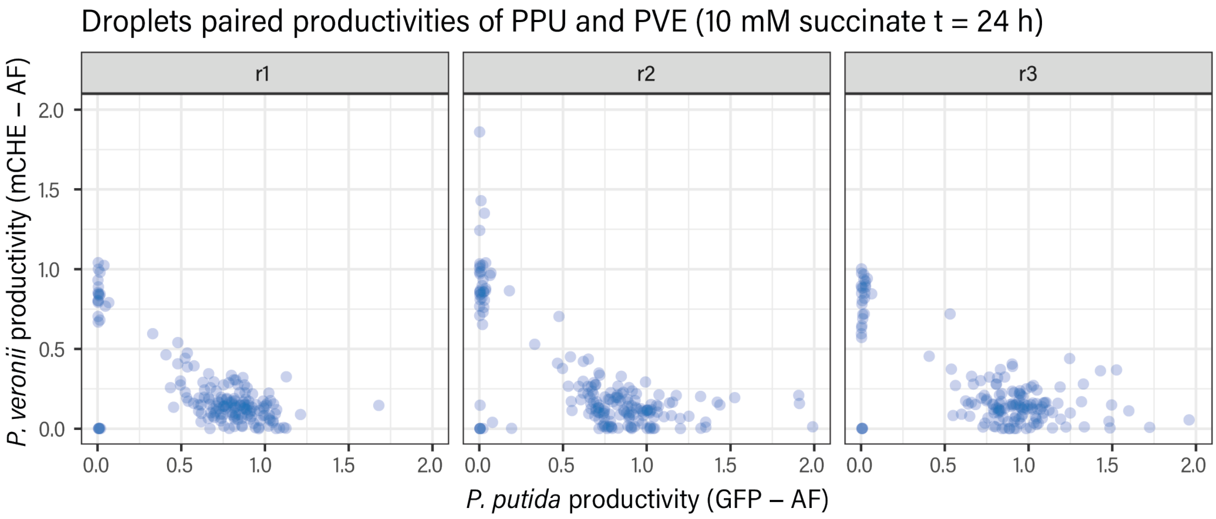


**Figure S12: Technical reproducibility of droplet cultivation procedures.** Paired productivities of *P. putida* (PPU) and *P. veronii* (PVE) after 24 h in a triplicate emulsion generated from a single suspension with 10 mM of succinate, to assess and demonstrate the technical reproducibility of droplet cultivation procedures.

**Supplementary movies:**

**Movie S1**: **Focal movie on one droplet where *P. protegens* CHA0 and Pf-5 are co-growing on MM C21 + 10 mM of succinate**. One picture/10 minutes is displayed.

**Movie S2: Focal movie on one droplet where *P. protegens* Pf-5 cells get all lysed while co-growing with CHA0 on MM C21 + 10 mM of succinate.** One picture/10 minutes is displayed.
